# Heterologous expression of microbial nitroreductases for TNT degradation in transgenic animals

**DOI:** 10.1101/2025.06.11.659045

**Authors:** Dimitri Perusse, Kate Tepper, Adam Sychla, Samuel J. Beach, Michael J. Smanski, Maciej Maselko

## Abstract

TNT (2,4,6-trinitrotoluene) from unexploded ordinances is a common environmental pollutant near munitions factories, military training sites, and areas of armed conflict. As physical and chemical approaches for TNT remediation are costly and difficult to scale, *in situ* bioremediation is an attractive alternative. TNT is a phytoaccumulative pollutant. Herbivores engineered to detoxify TNT are a potentially cost-effective platform to bioremediate large areas of contaminated sites while grazing or browsing. As a first step towards engineering herbivores with metabolic capabilities for TNT degradation, we engineered the animal genetic model, *Drosophila melanogaster*, to screen a library of microbial nitroreductases that can catalyse the initial degradation steps required for TNT degradation. We find strong, cofactor-dependent activity in fly lysates engineered to express NsfA from *Escherichia coli*, and NsfI from *Enterobacter cloacae*.

## Introduction

Pollution from explosives, including TNT, is a widespread problem. According to the Food and Agriculture Organisation of the United Nations, very high levels of pollution from explosives occurs in Southeast Asia, the Middle East, and Africa^1^. Investigative journalism by Propublica in 2017 following freedom of information requests revealed that the US is estimated to spend a billion dollars a year to remediate >40,000 sites that are contaminated with explosives^2^. Explosives used in the current wars in Ukraine and Gaza are expected to leave behind legacies of environmental pollution, which include TNT contamination^3–5^. Ukraine is one of the largest exporters of grains in the world, and as of March 2023, over a quarter of their agricultural land is estimated to be contaminated with explosives^3,4^.

TNT released into the environment is highly recalcitrant to degradation. It can adsorb to the soil matrix and leach into groundwater, where it can persist for many decades to centuries^6^. Some sites contaminated during World War II still present high concentrations of TNT^7^. Natural nitroaromatic compounds are rare, which may explain why TNT is not readily mineralized biologically^8^.

TNT toxicity can have broad impacts. In animals, TNT causes hemolysis leading to anaemia and damages blood conditioning organs such the liver, kidneys, and spleen^9,10^. Rats exposed to TNT over 2 years developed bladder carcinomas^9^, and it is toxic to the male reproductive system^9,10^. In plants, it damages the root system, stunts growth, and reduces seedling emergence^11–13^. It is also toxic to organisms important for soil health, such as microorganisms, fungi, and meso- and macro-organisms^12,14^. In aquatic environments, TNT can impede algae growth, and reduce survival of invertebrates, amphibians, and fish^13^.

A common method to remediate polluted soils is through composting^8^. This process relies on the ability of some microorganisms to degrade TNT into less toxic molecules^8,15^. TNT in contaminated soil may also be mineralized into its inorganic chemical constituents by incineration^8,16,17^. Groundwater is usually treated by filtration over granular activated carbon, or by Fenton processes^16,17^. All these processes can be expensive, requiring dedicated infrastructure, or a large-scale and disruptive reshaping of the environment. The US is reported to have spent over $40 billion US dollars on remediating pollution from explosives, and is expecting to pay a further $28 billion US dollars to complete the clean-up^2^.

Bioremediation is an attractive economical solution to clean up TNT from large areas with low-concentrations, and where the clean-up of a contaminated site can take a long time. Many plants accumulate TNT from the soil when they grow on contaminated sites^18^. Even if they possess the metabolic capacity to metabolize this chemical^19^, the process is slow and does not complete the break-down of TNT ^20,21^. However, several enzymes have been characterized that rapidly metabolize TNT^22^. This led French *et al*., almost twenty-five years ago to engineer tobacco plants to express PETN reductase from the bacterium *Enterobacter cloacae* PB2 for phytoremediation^23^. PETN reductase catalyses the removal of one nitro group from TNT^24^, allowing it to be conjugated to glutathione, which facilitates its sequestration into vacuoles or the lignin fraction of cell walls^25^. Following this, multiple enzymes from bacteria or animals have been introduced in different plants to confer the ability to metabolize multiple nitroaromatic pollutants^26–31^.

Although phytoremediation remains attractive, its deployment at scale is challenging. Phytoremediation requires first removing the native plants present in a contaminated site and then growing the transgenic phytoremediator over many years^28^. Following clean up, the transgenic plants are removed before attempting to re-establish the native flora. Biocontainment strategies need to be robust to avoid the undesired spread of transgenic plants in the area^32^.

A more scalable approach would involve engineering large herbivores to break down explosives, including TNT, in their diet (Figure 1C). These animals can be easily transported between contaminated sites and they would metabolize the TNT that accumulates in native flora, as they naturally break down plant matter during digestion. Biocontainment of fewer and larger transgenic animals is likely to be more practically achieved relative to phytoremediation approaches by fencing, surgical sterilization, or genetic approaches^33^ until TNT levels have dropped below an appropriate threshold.

**Figure 1:**
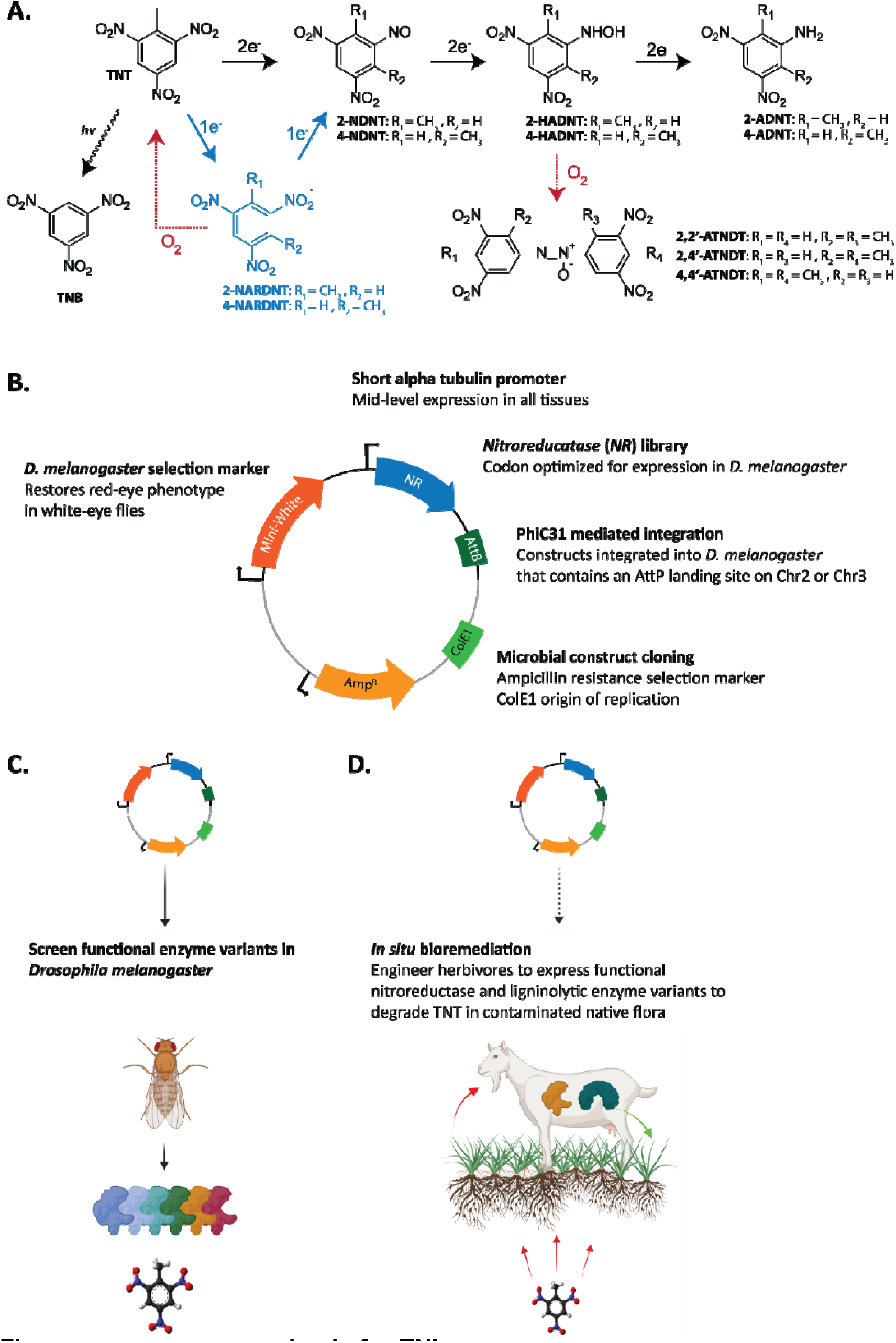
Engineering animals for TNT bioremediation. A. Chemical transformations involved in TNT bioremediation. Blue arrows and intermediates denote the oxygen-sensitive pathway. Red arrows show spontaneous oxidations. B. Diagram of the genetic constructs used to engineer *D. melanogaster* to express microbial nitroreductases. C. Functional microbial nitroreductases are first screened using *D. melanogaster*. D. Herbivores, such as goats and sheep, could subsequently be engineered to detoxify explosives, such as TNT, that accumulate in plants growing on contaminated sites. TNT: 2,4,6-Trinitrotoluene, TNB: 1,3,5-Trinitrobenzene, NARDNT: Nitroanion radical-dinitrotoluene, NDNT: Nitroso-dinitrotoluene, HADNT: Hydrolamino-dinitrotoluene, ATNDT: Azoxy-tetranitroditoluene, ADNT: Amino-dinitrotoluene.

Enzymatic approaches for TNT detoxification involve first the removal of the nitro groups or their transformation to facilitate further degradation reactions^8,25^. This can be achieved by microbial nitroreductases, which use electrons from NADH or NADPH to catalyse the stepwise reduction of one or more of the nitro-groups of TNT into nitroso-, then hydroxylamino-, and finally amino-group(s) (Figure 1A)^8^. Though these breakdown products and their condensation products, azoxynitrotoluenes, remain toxic, reduction of one or more of the nitro groups allows the subsequent oxidation of aromatic ring of TNT^8^. This oxidation step can be achieved using various ligninolytic oxidative enzymes from white-rot fungi, to complete the mineralisation of TNT^8^. Though ligninolytic enzymes would be toxic to plants and the biocontainment would be challenging in microbes, animals can be engineered to express functional ligninolytic enzymes to oxidize various aromatic industrial pollutants^34^. Functional variants of nitroreductases could be co-expressed with ligninolytic enzymes in an animal host to complete the detoxification of TNT.

As a first step toward herbivore TNT bioremediation, we engineered the animal model *D. melanogaster* to express six nitroreductases and characterized their activity *in vitro*. The functional microbial nitroreductases may provide a selection of low risk candidates that can be engineered into long lived herbivores (Figure 1).

## Results

### Engineering flies to express a library of microbial nitroreductases

Six bacterial nitroreductases previously described in the literature were engineered into *D. melanogaster* (Table 1). These included: *nsfI from E. cloacae 96-3*^37^, *nfsA from E. coli*^38^, *pnrA* from *Pseudomonas putida* JRL11^39^, *snrA from Salmonella enterica serovar Typhimurium* TA1535^40^, *mm-nr from Morganella morganii B2*^41^, and *xenA from P. putida* II-B^42^ (Table 1). Most of the enzymes are oxygen-insensitive nitroreductases. Each nitro group reduction requires two electrons, leading directly to the first nitroso intermediates 2-nitroso-4,6-dinitrotoluene or 4-nitroso-2,6-dinitrotoluene (Figure 1a)^37–41^. The enzyme XenA, on the other hand, is an oxygen-sensitive nitroreductase. In this case, the reduction of the nitro moiety into a nitroso is carried out by two successive one-electron transfers with the formation of a nitroanion radical intermediate^8^. This nitroanion radical can be easily quenched by a molecule of dioxygen to regenerate TNT (Figure 1). In most cases, the enzymes accept only NADPH as their electron source, except for NsfI, which can use NADPH and NADH to carry out the successive reductions^37–41^.

**Table 1:** Nitroreductase genes engineered into *D. melanogaster*.

| Gene name | Genbank | Enzyme | Native host | Prior heterologous expression in eukaryotes |
| --- | --- | --- | --- | --- |
| *nsfl* | AAA62801 | Oxygen-insensitive NAD(P)H nitroreductase | *E. cloacae* | Tobacco, switchgrass |
| *nsfA* | BAA07425 | Oxygen-insensitive NADPH nitroreductase | *E. coli* | *Arabidopsis thaliana*, human cell lines |
| *pnrA* | AAM95986 | Oxygen-insensitive NADPH nitroreductase | *P. putida* | *Populus tremula* x *P. tremuloides* |
| *XenA* | AF154061_1 | Oxygen-sensitive Xenobiotic reductase | *P. putida* II-B | *E. coli* |
| *snrA* | AAD18027 | Nitroreductase A homolog | *S. enterica* | N/A |
| *mm-nr* | AGG30128 | Oxygen-insensitive NAD(P)H nitroreductase | *M. morganii* | N/A |

Three of the six microbial nitroreductases have previously been expressed in a eukaryotic host. The genes *nsfI, nsfA*, and *pnrA* were engineered into *Arabidopsis thaliana*^43^, tobacco^29^, aspen^44^, or grasses^45^ for the purpose of TNT bioremediation. Transgenic human cell lines involving *nsfA* were created to activate prodrugs containing nitro groups into cytotoxic chemicals for cancer research^46^.

The nitroreductases open reding frames were codon optimised for *D. melanogaster* and expressed from a truncated alpha tubulin promoter (Figure 1B). We selected this promoter as it has been used previously to heterologously express enzymes for prototyping animal bioremediation in flies^34,47^. It provides expression likely across a broad set of tissues, and at a moderate level to reduce potential toxic overexpression^48^. The expression vectors were integrated into *D. melanogaster* strains harboring AttP landing sites using ϕC31 mediated integration^49^. Transgenic *D. melanogaster* were selected using the *mini-white marker*^50^. We validated the genomic integration of the expression vectors by PCR and Sanger sequencing (data not shown). We also validated nitroreductase mRNA expression from the truncated alpha tubulin promoter by RT-qPCR. The nitroreductases’ mRNA levels were between 10% and 30% of the housekeeping reference gene EF1 (Supplementary Figure 1).

### *In vitro* TNT degradation

We tested the library of microbial nitroreductases engineered into *D. melanogaster* for their ability to degrade TNT *in vitro*. Triplicates of transgenic fly lysates and w^1118^ fly lysate controls were incubated with 50 µM TNT and 300 µM NADH or NADPH as an electron donor for 90 mins. TNT concentrations were measured by HPLC.

With NADH as the electron donor, only the fly lysates prepared from adult flies expressing NsfI were able to degrade TNT (Figure 2A). At the 90 min timepoint, 5.5% of TNT remained in solution. No significant degradation activity was detected for the other fly lysates compared to the w^1118^ control within the allocated experimental time (Figure 2B).

**Figure 2:**
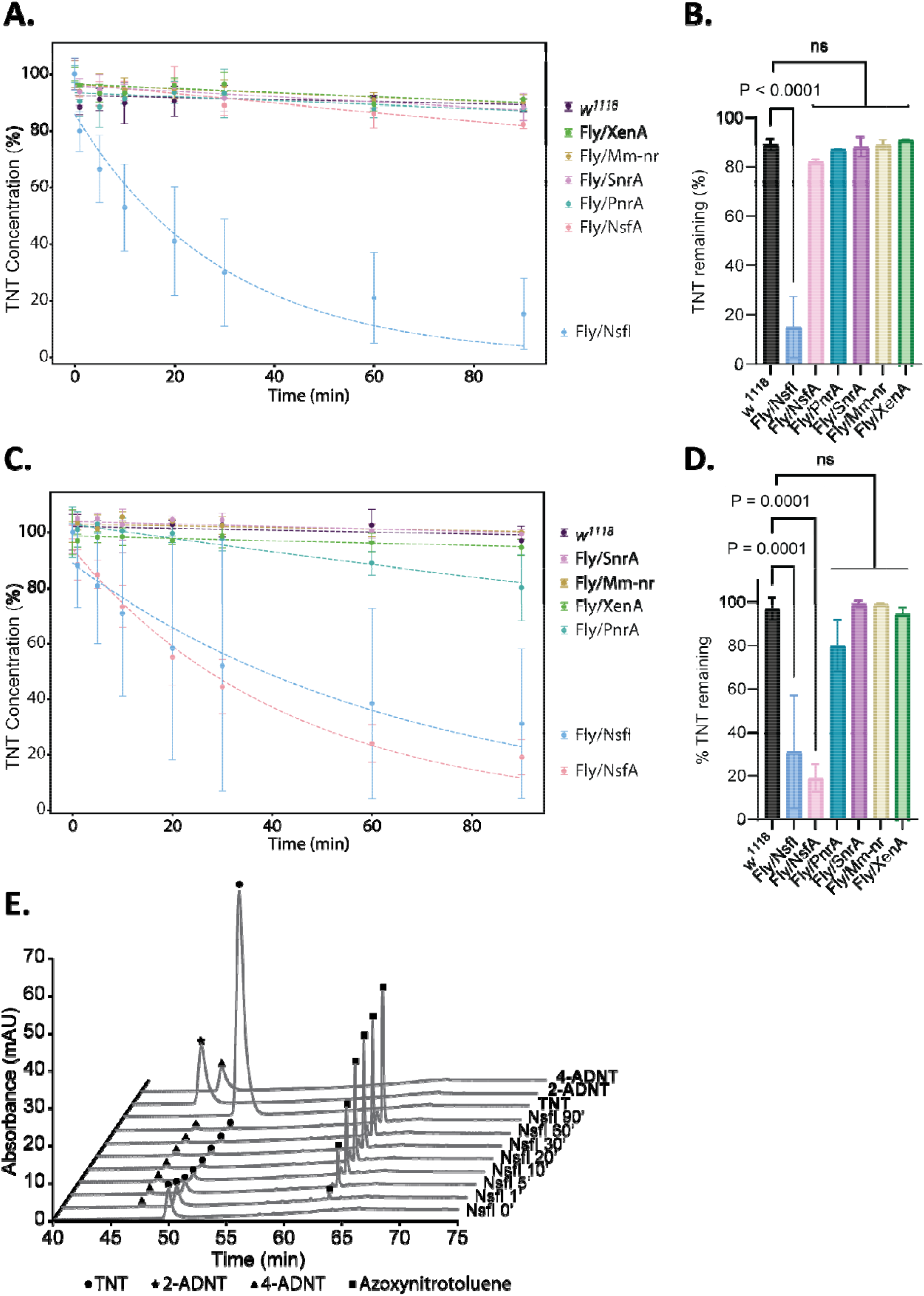
TNT degradation in fly lysates. Fly lysates and controls were incubated with 50 µM TNT with 300 µM **A**. NADH or **C**. NADPH in 25 mM Tris-HCl pH 7.4 for 90 minutes. TNT concentrations were measured using HPLC. *n* = 3 biologically independent replicates. Statistical analysis of **A**. and **C**. were performed at end-point in **B**. and **D**., respectively. The statistical analysis was conducted using one-way ANOVA using Dunnett’s method compared to controls, where ns = not statistically significant (p > 0.05). E. HPLC chromatograms for NsfI fly lysates incubated with 300 µM NADH and 50 µM TNT over a 90-minute time course in comparison to pure standards for TNT and its putative breakdown products 2-ADNT and 4-ADNT. w^1118^ = *D. melanogaster* genetic background for the engineered strains, Fly/NsfA = *D. melanogaster* engineered to express *nsfA* from *E. coli*, Fly/PnrA = *D. melanogaster* engineered to express *pnrA* from *P. putida*, Fly/SnrA = *D. melanogaster* engineered to express *snrA* from *S. enterica*, Fly/Mm-nr = *D. melanogaster* engineered to express a nitroreductase from *M. morganii*, and Fly/NsfI = *D. melanogaster* engineered to express *nsfI* from *E. cloacae*.

According to previous studies^38–42^, most of the enzymes selected for this experiment use NADPH as the electron donor to reduce the nitro moieties of TNT. Using NADPH as an electron donor, we observed degradation of TNT in extracts from flies expressing NsfA and NsfI, with 19% and 30% of the TNT remaining after 90 min, respectively (Figure 2C). The difference between degradation by NsfA/I and the w^1118^ control was statistically significant (Figure 2D). NsfA and NsfI were not statistically different from each other (Figure 2D). Fly lysates prepared from flies expressing PnrA show a slight decrease in TNT concentration in later timepoints (Figure 2C), but the result is not statistically significant compared to the w^1118^ control (Figure 2D). The other enzymes, SnrA, Mm-nr, and XenA, and the w^1118^ control did not show TNT degradation within the allocated experimental time (Figure 2C).

### TNT degradation products

The HPLC chromatogram peaks corresponding to TNT and its expected amino-dinitrotoluene degradation products were annotated by comparison to standards (Figure 2E). We observed a small peak corresponding to 4-amino-dinitrotoluene (4-ADNT), though it was not equivalent to the initial amount of TNT. We observed a small and large peak of uncharacterized compounds with a retention time of 64 minutes concomitant with TNT degradation. We postulate that these are isomers of azoxynitrotoluene degradation products (Figure 1A). According to the degradation pathway, the hydroxylamino intermediates can spontaneously dimerize in oxygenated conditions, particularly *in vitro*, to form azoxynitrotoluene molecules (Figure 1A)^8^. We did not confirm this identity by NMR or spectroscopic analysis and do not have authentic standards for the azoxynitrotoluene compounds.

## Discussion

This research is the first step towards engineering long lived herbivores, such as goats and sheep, to bioremediate large areas of phytoaccumulative pollutants, such as TNT. Using *D. melanogaster* as a research tool to screen a library of nitroreductases, we identified NsfI and NsfA are capable of catalysing the first step of TNT degradation *in vitro* to remove the electron withdrawing nitro-groups of TNT. These enzymes could be combined with the ligninolytic enzymes we characterised previously in *D. melanogaster* to oxidize the TNT intermediates and complete the detoxification of TNT. Future experiments will involve engineering this enzyme pathway into short lived herbivores, such as rabbits, to prototype animal TNT bioremediation from contaminated plant material and to characterise the breakdown products of the enzyme pathway.

Herbivore bioremediation could add unique capabilities to the field of bioremediation. The herbivores can be readily transported between sites and immediately process contaminated native plants. Robust transgene biocontainment can be achieved in herbivores such as goats and sheep, using industry standard surgical sterilization. The animals may also be fenced into specific areas, and/or tracked via GPS. Animals may be engineered to express ligninolytic oxidative enzymes to complete the detoxification of TNT. Though microbes commonly metabolize TNT to the toxic 2-ADNT and 4-ADNT intermediates in aerobic soils, engineering microbes to express oxidative enzymes from white-rot fungi is limited by effective transgene biocontainment strategies required for their wild release^51^. Engineering plants to express ligninolytic oxidative enzymes is challenging as these enzymes will also degrade lignin in plants^52^.

A major disadvantage of using herbivores for TNT bioremediation is that TNT predominantly accumulates in the root material, with less than 25% of TNT in the aerial parts of the plant, depending on the species^18^. The TNT accessible to the herbivores will therefore be low, and would significantly increase the time it takes to remediate a site. Another major disadvantage is that any pollutant present in plant material is in equilibrium with the soil, and to remediate a pollutant from the soil will also take a long time^28^. These disadvantages may be acceptable in large contaminated areas where no other economical remediation option is available and time is not an important factor. Where duration is an important factor, it could be addressed by combining herbivore bioremediation with phytoremediators engineered to hyperaccumulate TNT into the aerial parts of the plant.

Previously we showed that animals engineered to express mercury detoxification enzymes were resistant to greater methylmercury exposure^47^. We expect that animals engineered for TNT detoxification will be similarly resistant to greater TNT exposure. They may therefore remediate a site for wildlife that may be harmed by TNT in the contaminated area, without harm to themselves. Future studies to determine TNT toxicity in wildtype and engineered herbivores will be critical.

## Materials and Methods

### Fly expression plasmid assembly

The nitroreductase library enzyme sequences (Table 1) were codon optimized (Integrated DNA Technologies (IDT) codon optimization tool) for expression in D. melanogaster. The *D. melanogaster* Kozak consensus sequence: “AATCTTACAAA” was added immediately preceding the start codon. At least 20 bp of homology arms to the destination plasmid (pMC-1-1-1)^34^ were added upstream and downstream. The sequences were ordered as gBlocks from IDT and assembled (NEBuilder® HiFi DNA assembly master mix (New England Biolabs (NEB))) into the pMC-1-1-1 expression plasmid linearized with NotI-HF. The plasmids were verified by Sanger sequencing. See Supplementary Table S3 and S4 for plasmid descriptions and sequences, and Supplementary Table S2 for primers used in this study.

### *D. melanogaster* husbandry, strains, and transgenesis

All lines were maintained at 25°C and 12 h light/dark cycle and 60–70% relative humidity. The flies were kept in vials containing standard cornmeal/agar diet Bloomington formula (Genesee Scientific, 66–121), 0.05 M propionic acid (Sigma, 402907), and 0.1% tegosept (Genesee Scientific, 20–258).

w^1118^ *D. melanogaster* strains were used to control for genetic background. w^1118^ are WT flies that have the *mini-white* gene knocked out, resulting in white eyes.

The plasmids containing the different nitroreductases were sent to Bestgene Inc. (Chino Hills, CA) for ϕC31-mediated integration and outcrosses to balancer strains. Homozygous fly lines for the nitroreductase constructs were generated by selecting against balancer phenotypic markers. Homozygous NfsA flies could not be generated and heterozygous flies were used in the assays. See Supplementary Table S1 for fly strains used in this study.

### Validation of transgene genomic integration

Five frozen adult flies were homogenized using a motorized pestle in 500 µL 10 mM Tris-HCl pH 8 (Roche, #10812846001), 1 mM ethylenediaminetetraacetate (EDTA) (Millipore Sigma, #ED-500G), 25 mM sodium chloride (Macron Fine Chemicals, #7581-06), and 200 g/mL Proteinase K (New England Biolabs, #P8107S). The mix was then incubated at 37°C for 30 min, followed by 3 min at 95°C to inactivate the Proteinase K. The tube was centrifuged at 17,900 rcf for 5 min and 0.5 μL of the supernatant was collected for PCR analysis. The gDNA in solution was used as the template with a universal set of primers flanking the nitroreductase locus (Supplementary Table S2) and the region of interest was amplified with CloneAmpTM Hi-Fi PCR Premix (Takara Bio USA, #639298). The DNA thermocycling conditions were: initial denaturation step at 98°C for 3 min; 35 cycles of 98°C for 10 s, 70°C for 15 s, and 72°C for 30 s; and a final extension step at 72°C for 5 min. The PCR products were analyzed by agarose gel electrophoresis and Sanger sequencing (ACGT, Inc.).

### Validation of transgene mRNA expression

RNA was extracted from the flies using the TRIzol reagent (Thermo Fisher Scientific, #15596018) following the manufacturer instructions. Three frozen adult flies from the same line were homogenized with a motorized pestle in 0.2 mL TRIzol. An additional 0.8 mL of TRIzol was added. The RNA isolated was resuspended in nuclease-free water (Takara Bio USA, #9012) and then treated with DNase using the TURBO DNA-free kit (Invitrogen, #AM2238).

RT-qPCR was carried out from the isolated RNA (100 ng) using the Luna Universal One-Step RT-qPCR Kit (New England Biolabs, #E3005L) in a 12 μL reaction total volume following the manufacturer instructions. Fold changes for RT-qPCR were determined by the ΔΔCT method^35^. Expression of the nitroreductases were normalized to *D. melanogaster* gene Elongation Factor 1 alpha100 (EF1)^36^. See Supplementary Table S2 for primers used in this study.

### Chemicals and stock solution preparation

A slurry of 30% TNT in water was obtained from the US department of defence. A portion of the 30% TNT slurry was frozen at -80°C overnight and was lyophilized to remove the water from the sample. A 0.5 mM TNT solution was prepared in 25 mM Tris buffer solution at pH 7.4.

A TNT analytical standard was purchased from Millipore Sigma (#ERT-022S-1.2ML). Analytical standards for 2-amino-dinitrotoluene (2-ADNT) and 4-amino-dinitrotoluene (4-ADNT) were purchased from Millipore Sigma (#ERA-017-100MG and #ERA-018-100MG, respectively). Stock solutions of 2-DANT and 4-DANT (1 mg/mL) were prepared by dissolving each compound in acetonitrile (Fisher Scientific, A998-4). Serial dilutions were prepared from the stock solutions at 100, 50, 25, 10, 5, 2.5, 1, 0.5, and 0.1 µg/mL. See Supplementary Figures S2-S5 for the TNT, 2-ADNT, and 4-ADNT HPLC calibration curves.

Solutions of 3 mM NADH and NADPH (Millipore Sigma, #481913-500MG and Neta Scientific, #RPI-N20120-0.1, respectively) were made in 25 mM tris buffer solution at pH 7.4. The solutions were prepared immediately before experiments and kept on ice.

### *In vitro* degradation of TNT by adult fly lysates

To prepare fly lysate, 20 frozen adult flies were homogenized with a motorized pestle in 450 µL of 50 mM tris-HCl buffer pH 7.4. Each tube was incubated for 15 min at room temperature. Each tube was centrifuged at 17,000 rcf at room temperature for 10 min, and 350 μL of the supernatant was transferred into a clean microcentrifuge tube. The solution was centrifuged again at 17,000 rcf at room temperature for 10 min and 275 μL of supernatant was transferred into another clean tube. After a third centrifuge at the same speed, 200 μL of supernatant was transferred to another tube and diluted up to 1 mL in 25 mM Tris buffer pH 7.4.

To prepare the *in vitro* enzyme assay, 720 μL of the fly lysate prepared above was added to 90 μL of the 0.5 mM TNT stock solution. Then 90 μL aliquots of the enzyme-TNT solution were added into an 8-strip PCR tubes (Genesee, 27-426F). To start the enzymatic reaction, 10 μL of either 3 mM NADH or NADPH stock solutions was added. In the negative control, the electron donor solution was replaced by 10 μL of 25 mM Trizma buffer pH 7.4. The reaction mixes were incubated at room temperature for various durations as described in the results. To stop the reactions, the strip tube was incubated at 80°C for 5 min.

### TNT degradation analysis by high performance liquid chromatography (HPLC)

Samples (100 µL) were introduced into inserts (VWR, #97051-402) for 2 mL HPLC vials (ChromTech, #CTV-1209GSA). The HPLC analyses were performed on an Ultimate 3000 Rapid Separation system using either a Phenomenex Luna C18 column (4.6 × 150 mm, 3 µm) (#00F-4114-E0) or an Agilent Eclipse XDB-C18 column (4.6 × 100 mm, 3.5 µm) (#961967-902). The sample injection volume was 10 µL. All chromatography was performed at a flow rate of 1.2 mL/min using Milli-Q water filtered on 0.22 µm filter (Millipore Sigma, JGWP04700) (solvent A) and acetonitrile (Fisher Scientific, A998-4) (solvent B) with the different gradients depending on the column. Gradient for the Luna column: 0-65 min, 10% to 90% solvent B; 65 - 75 min, 95% solvent B; 77 - 81 min, 90% to 10% solvent B. Gradient for the Eclipse column: 0 - 10 min, 10% to 30% solvent B; 10 - 12.5 min, 30% solvent B; 12.5 - 25 min, 30% to 95% solvent B; 25 - 27 min, 95% solvent B, 27 - 28 min, 95% to 10% solvent B, 28 - 30 min 10% solvent B. Elution compounds were detected at λ = 254 nm.

## Figures

Figures were made using GraphPad Prism 10, Excel, Python, Biorender, and Adobe Illustrator.

## Acknowledgements

This research was funded by the International Technology Center Pacific (ITC-PAC) under Contract No. FA520923C0014 (M.M.). D.P. was supported by a fellowship from the MnDRIVE at the University of Minnesota. K.T. was supported by an Australian Government Research Training Program Scholarship.

The authors declare no competing financial interests.

## Author Contributions

K.T. and M.M. conceived of the study. K.T., D.P., and S.B. performed the experiments. All authors contributed to experimental designs and data analysis. All authors contributed to writing the manuscript. Co-first authors DP and KT have editorial approval to list their name in the first positions on their respective CVs.

## References

1. FAO and UNEP. Global Assessment of Soil Pollution: Report. 10.4060/cb4894en (2021).

2. Groeger, L., Grochowski Jones, R. & Lustgarten, A. Bombs in Your Backyard. Propublica https://projects.propublica.org/bombs/ (2017).

3. Nickel, R. Insight: Soils of war: The toxic legacy for Ukraine’s breadbasket. Reuters (2023).

4. Turns, A. Russia’s invasion is leaving a toxic trace on Ukrainian soil, contaminating crops and posing a serious long-term risk to human health. BBC https://www.bbc.com/future/article/20230221-the-toxic-legacy-of-the-ukraine-war (2023).

5. Webber, P. & Parkinson, S. Gaza: one of the most intense bombardments in history? SGR https://www.sgr.org.uk/resources/gaza-one-most-intense-bombardments-history (2023).

6. Amaral, H. I. F., Fernandes, J., Berg, M., Schwarzenbach, R. P. & Kipfer, R. Assessing TNT and DNT groundwater contamination by compound-specific isotope analysis and 3H–3He groundwater dating: A case study in Portugal. Chemosphere 77, 805–812 (2009).

7. Eisentraeger, A., Reifferscheid, G., Dardenne, F., Blust, R. & Schofer, A. Hazard characterization and identification of a former ammunition site using microarrays, bioassays, and chemical analysis. Environ Toxicol Chem 26, 634–646 (2007).

8. Esteve-Núñez, A., Caballero, A. & Ramos, J. L. Biological Degradation of 2,4,6-Trinitrotoluene. Microbiology and Molecular Biology Reviews 65, 335–352 (2001).

9. Johnson, M. S. & Reddy, G. Chapter 3 - Wildlife Toxicity Assessment for 2,4,6-Trinitrotoluene (TNT). in Wildlife Toxicity Assessments for Chemicals of Military Concern (eds. Williams, M. A., Reddy, G., Quinn, M. J. & Johnson, M. S.) 25–51 (Elsevier, 2015). doi:10.1016/B978-0-12-800020-5.00003-X.

10. Agency for Toxic Substances and Disease Registry. TOXICOLOGICAL PROFILE FOR 2,4,6-TRINITROTOLUENE. (1995).

11. Simini, M. et al. Toxicities of TNT and RDX to Terrestrial Plants in Five Soils with Contrasting Characteristics. U.S. Army Research, Development and Engineering Command (2013).

12. Best, E. P. H., Tätern, H. E., Geter, K. N., Wells, M. L. & Lane, B. K. Toxicity and Metabolites of 2,4,6-Trinitrotoluene (TNT) in Plants and Worms from Exposure to Aged Soil. (2004).

13. Lotufo, G. R. Toxicity and Bioaccumulation of Munitions Constituents in Aquatic and Terrestrial Organisms. in Energetic Materials: From Cradle to Grave (eds. Shukla, M. K., Boddu, V. M., Steevens, J. A., Damavarapu, R. & Leszczynski, J.) 445–479 (Springer International Publishing, Cham, 2017). doi:10.1007/978-3-319-59208-4_13.

14. Meyers, S. K., Deng, S., Basta, N. T., Clarkson, W. W. & Wilber, G. G. Long-Term Explosive Contamination in Soil: Effects on Soil Microbial Community and Bioremediation. Soil and Sediment Contamination: An International Journal 16, 61–77 (2007).

15. Rezaei, M. R., Abdoli, M. A., Karbassi, A., Baghvand, A. & Khalilzadeh, R. Bioremediation of TNT Contaminated Soil by Composting with Municipal Solid Wastes. Soil and Sediment Contamination: An International Journal 19, 504–514 (2010).

16. US EPA. Technical Fact Sheet – 2,4,6-Trinitrotoluene (TNT). (2014).

17. US EPA. Technical Fact Sheet – 2,4,6-Trinitrotoluene (TNT). (2017).

18. Vila, M., Lorber-Pascal, S. & Laurent, F. Fate of RDX and TNT in agronomic plants. Environmental Pollution 148, 148–154 (2007).

19. Gandia-Herrero, F. et al. Detoxification of the explosive 2,4,6-trinitrotoluene in Arabidopsis: discovery of bifunctional O- and C-glucosyltransferases. The Plant Journal 56, 963–974 (2008).

20. Rylott, E. L., Lorenz, A. & Bruce, N. C. Biodegradation and biotransformation of explosives. Curr Opin Biotechnol 22, 434–440 (2011).

21. Adamia, G. et al. Absorption, distribution, and transformation of TNT in higher plants. Ecotoxicol Environ Saf 64, 136–145 (2006).

22. Roldán, M. D., Pérez-Reinado, E., Castillo, F. & Moreno-Vivián, C. Reduction of polynitroaromatic compounds: the bacterial nitroreductases. FEMS Microbiol Rev 32, 474–500 (2008).

23. French, C. E., Rosser, S. J., Davies, G. J., Nicklin, S. & Bruce, N. C. Biodegradation of explosives by transgenic plants expressing pentaerythritol tetranitrate reductase. Nat Biotechnol 17, 491–494 (1999).

24. French, C. E., Nicklin, S. & Bruce, N. C. Aerobic Degradation of 2,4,6-Trinitrotoluene by Enterobacter cloacae PB2 and by Pentaerythritol Tetranitrate Reductase. Appl Environ Microbiol 64, 2864–2868 (1998).

25. Rylott, E. L. & Bruce, N. C. Right on target: using plants and microbes to remediate explosives. Int J Phytoremediation 21, 1051–1064 (2019).

26. Rylott, E. L. et al. An Explosive-Degrading Cytochrome P450 Activity and Its Targeted Application for the Phytoremediation of RDX. Nature Biotechnology vol. 24 10.1038/nbt1184 (2006).

27. Zhang, L., Rylott, E. L., Bruce, N. C. & Strand, S. E. Genetic modification of western wheatgrass (Pascopyrum smithii) for the phytoremediation of RDX and TNT. Planta 249, 1007–1015 (2019).

28. Cary, T. J. et al. Field trial demonstrating phytoremediation of the military explosive RDX by XplA/XplB-expressing switchgrass. Nat Biotechnol 39, 1216–1219 (2021).

29. Hannink, N. K. et al. Enhanced Transformation of TNT by Tobacco Plants Expressing a Bacterial Nitroreductase. Int J Phytoremediation 9, 385–401 (2007).

30. Hannink, N. K., Rosser, S. J., French, C. E. & Bruce, N. C. Uptake and Metabolism of TNT and GTN by Plants Expressing Bacterial Pentaerythritol Tetranitrate Reductase. Water, Air and Soil Pollution: Focus 3, 251–258 (2003).

31. van Dillewijn, P. et al. Bioremediation of 2,4,6-Trinitrotoluene by Bacterial Nitroreductase Expressing Transgenic Aspen. Environ Sci Technol 42, 7405–7410 (2008).

32. Clark, M. & Maselko, M. Transgene Biocontainment Strategies for Molecular Farming. Front Plant Sci 11, 210 (2020).

33. Maselko, M. et al. Engineering multiple species-like genetic incompatibilities in insects. Nat Commun 11, 1–7 (2020).

34. Clark, M. et al. Bioremediation of Industrial Pollutants by Insects Expressing a Fungal Laccase. ACS Synth Biol 11, 308–316 (2022).

35. Livak, K. J. & Schmittgen, T. D. Analysis of Relative Gene Expression Data Using Real-Time Quantitative PCR and the 2–ΔΔCT Method. Methods 25, 402–408 (2001).

36. Ponton, F., Chapuis, M.-P., Pernice, M., Sword, G. A. & Simpson, S. J. Evaluation of potential reference genes for reverse transcription-qPCR studies of physiological responses in Drosophila melanogaster. J Insect Physiol 57, 840–850 (2011).

37. Bryant, C. & DeLuca, M. Purification and characterization of an oxygen-insensitive NAD(P)H nitroreductase from Enterobacter cloacae*. Journal of Biological Chemistry 266, 4119–4125 (1991).

38. Zenno, S. et al. Biochemical characterization of NfsA, the Escherichia coli major nitroreductase exhibiting a high amino acid sequence homology to Frp, a Vibrio harveyi flavin oxidoreductase. J Bacteriol 178, 4508–4514 (1996).

39. Caballero, A., Lázaro, J. J., Ramos, J. L. & Esteve-Núñez, A. PnrA, a new nitroreductase-family enzyme in the TNT-degrading strain Pseudomonas putida JLR11. Environ Microbiol 7, 1211–1219 (2005).

40. Nokhbeh, M. R. et al. Identification and characterization of SnrA, an inducible oxygen-insensitive nitroreductase in Salmonella enterica serovar Typhimurium TA1535. Mutation Research/Fundamental and Molecular Mechanisms of Mutagenesis 508, 59–70 (2002).

41. Kitts, C. L., Green, C. E., Otley, R. A., Alvarez, M. A. & Unkefer, P. J. Type I nitroreductases in soil enterobacteria reduce TNT (2,4,6-trinitrotoluene) and RDX (hexahydro-1,3,5-trinitro-1,3,5-triazine). Can J Microbiol 46, 278–282 (2000).

42. Blehert, D. S., Fox, B. G. & Chambliss, G. H. Cloning and Sequence Analysis of Two PseudomonasFlavoprotein Xenobiotic Reductases. J Bacteriol 181, 6254–6263 (1999).

43. Kurumata, M. et al. Tolerance to, and Uptake and Degradation of 2,4,6-Trinitrotoluene (TNT) are Enhanced by the Expression of a Bacterial Nitroreductase Gene in Arabidopsis thaliana. 60, 272–278 (2005).

44. van Dillewijn, P. et al. Bioremediation of 2,4,6-Trinitrotoluene by Bacterial Nitroreductase Expressing Transgenic Aspen. Environ Sci Technol 42, 7405–7410 (2008).

45. Zhang, L. et al. Expression in grasses of multiple transgenes for degradation of munitions compounds on live-fire training ranges. Plant Biotechnol J 15, 624–633 (2017).

46. Vass, S. O., Jarrom, D., Wilson, W. R., Hyde, E. I. & Searle, P. F. E. coli NfsA: an alternative nitroreductase for prodrug activation gene therapy in combination with CB1954. Br J Cancer 100, 1903–1911 (2009).

47. Tepper, K. et al. Methylmercury demethylation and volatilization by animals expressing microbial enzymes. bioRxiv (2023) doi:10.1101/2023.12.11.571038.

48. Waters, A. J., Capriotti, P., Gaboriau, D. C. A., Papathanos, P. A. & Windbichler, N. Rationally-engineered reproductive barriers using CRISPR & CRISPRa: an evaluation of the synthetic species concept in Drosophila melanogaster. Sci Rep 8, 13125 (2018).

49. PhiC31 integrase-mediated transgenesis systems. Best Gene https://www.thebestgene.com/PhiC31InfoPage.do (2023).

50. Klemenz, R., Weber, U. & Gehring, W. J. The white gene as a marker in a new P-element vector for gene transfer in Drosophila. Nucleic Acids Res 15, 3947–3959 (1987).

51. George, D. R. et al. A bumpy road ahead for genetic biocontainment. Nat Commun 15, 650 (2024).

52. Plácido, J. & Capareda, S. Ligninolytic enzymes: a biotechnological alternative for bioethanol production. Bioresour Bioprocess 2, 23 (2015).

