## Supplemental File 1 for "Heterologous expression of microbial nitroreductases for TNT degradation in transgenic animals"

1. Department of Biochemistry, Molecular Biology, and Biophysics. University of Minnesota, Saint Paul, MN 55108
2. Biotechnology Institute, University of Minnesota, Saint Paul, MN 55108
3. Applied BioSciences, Macquarie University, Sydney, NSW 2109, Australia
4. Australian Research Council Centre of Excellence in Synthetic Biology, Macquarie University, Sydney, NSW 2109, Australia

*Joint first authors who may be interchangeably listed first

Supplementary Figures 1

Supplementary Tables 1-4

Supplementary References

**Supplementary Figure S1**

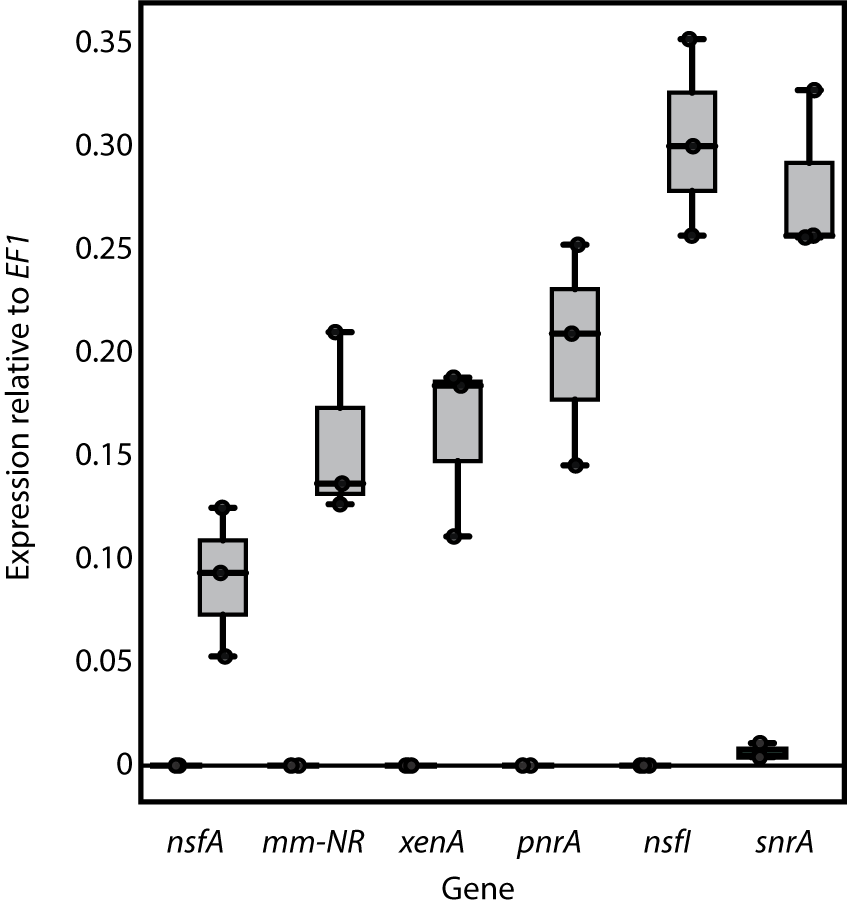

**Fig. S1. Expression of nitroreductase genes in transgenic *D. melanogaster*.**
Three biological replicates are reported for transgenic (right box plots) and WT (left box plots) for each nitroreductase gene listed on the X-axis.

**Supplementary figure S2**

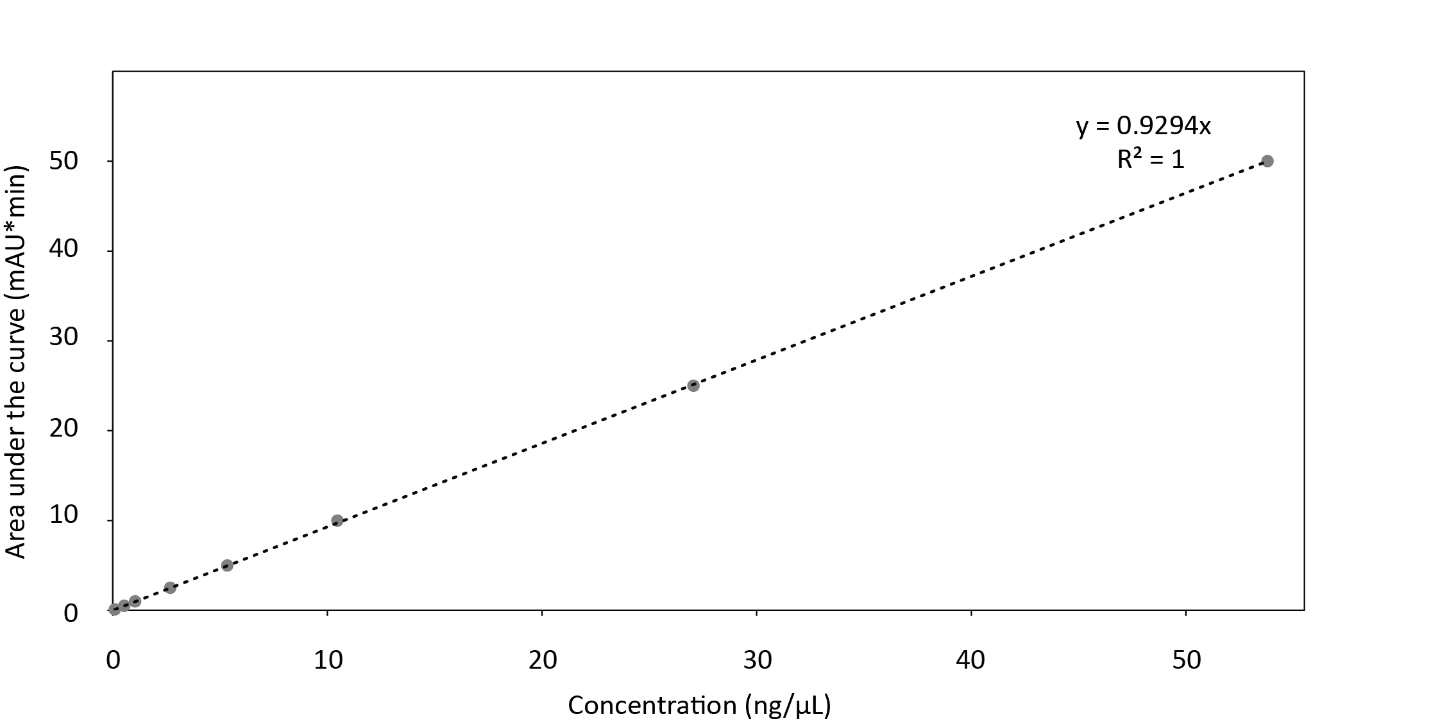

**Fig S2. 2,4,6-trinitrotoluene calibration curve with the Luna column.**

**Supplementary figure 3**

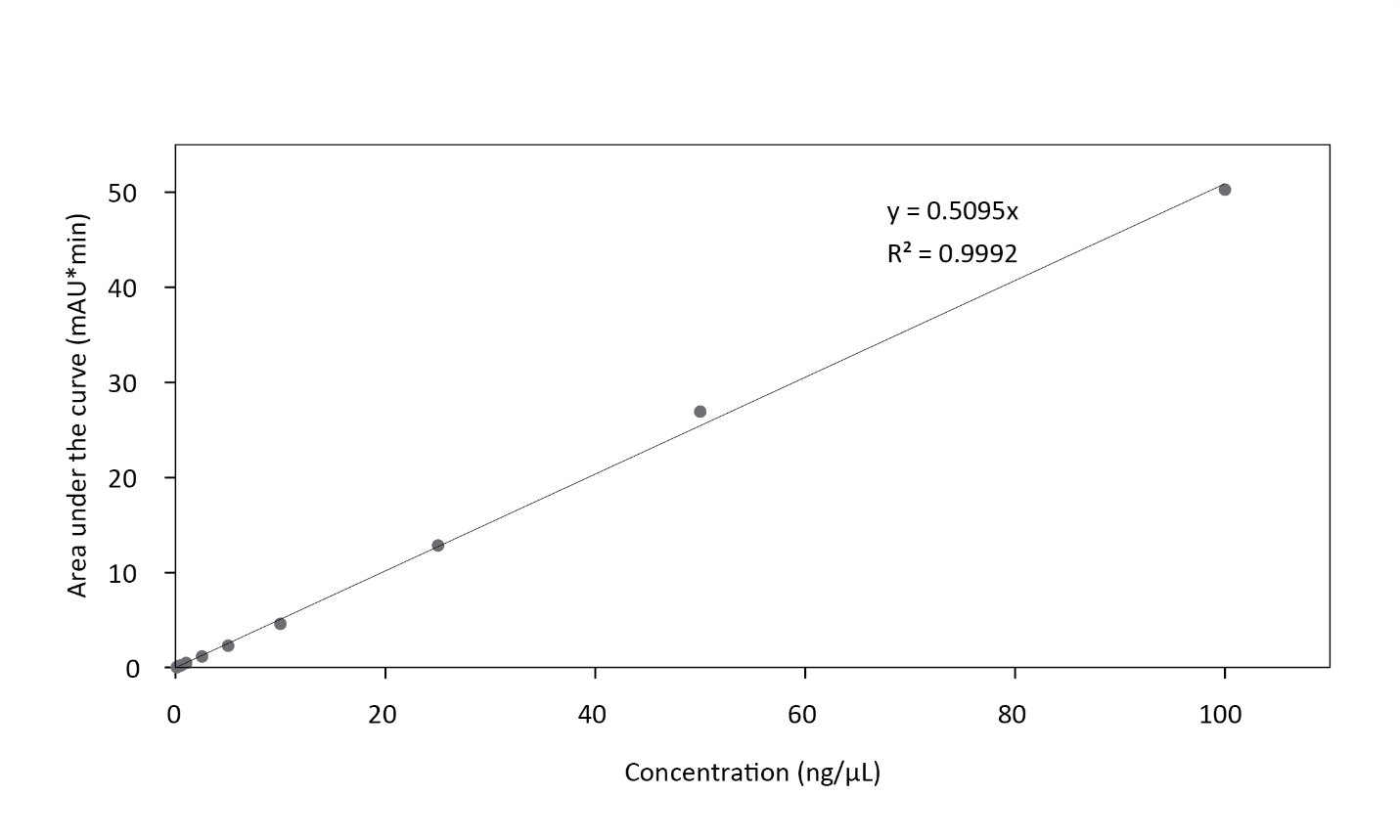

**Fig S3. 2,4,6-trinitrotoluene calibration curve with the Eclipse column.**

**Supplementary figure S4**

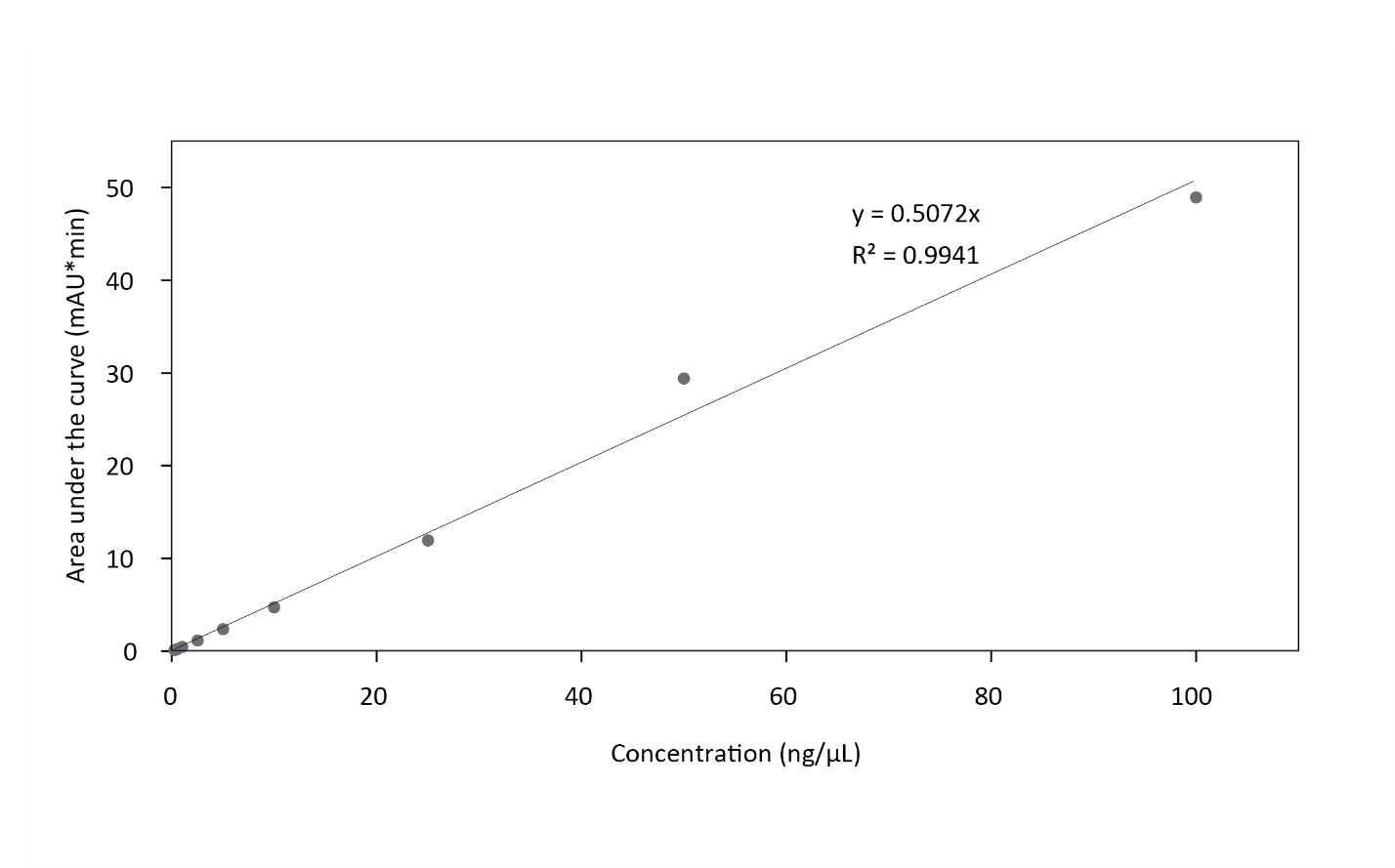

**Fig S4. 2-amino-4,6-dinitrotoluene calibration curve with the Eclipse column.**

**Supplementary figure S5**

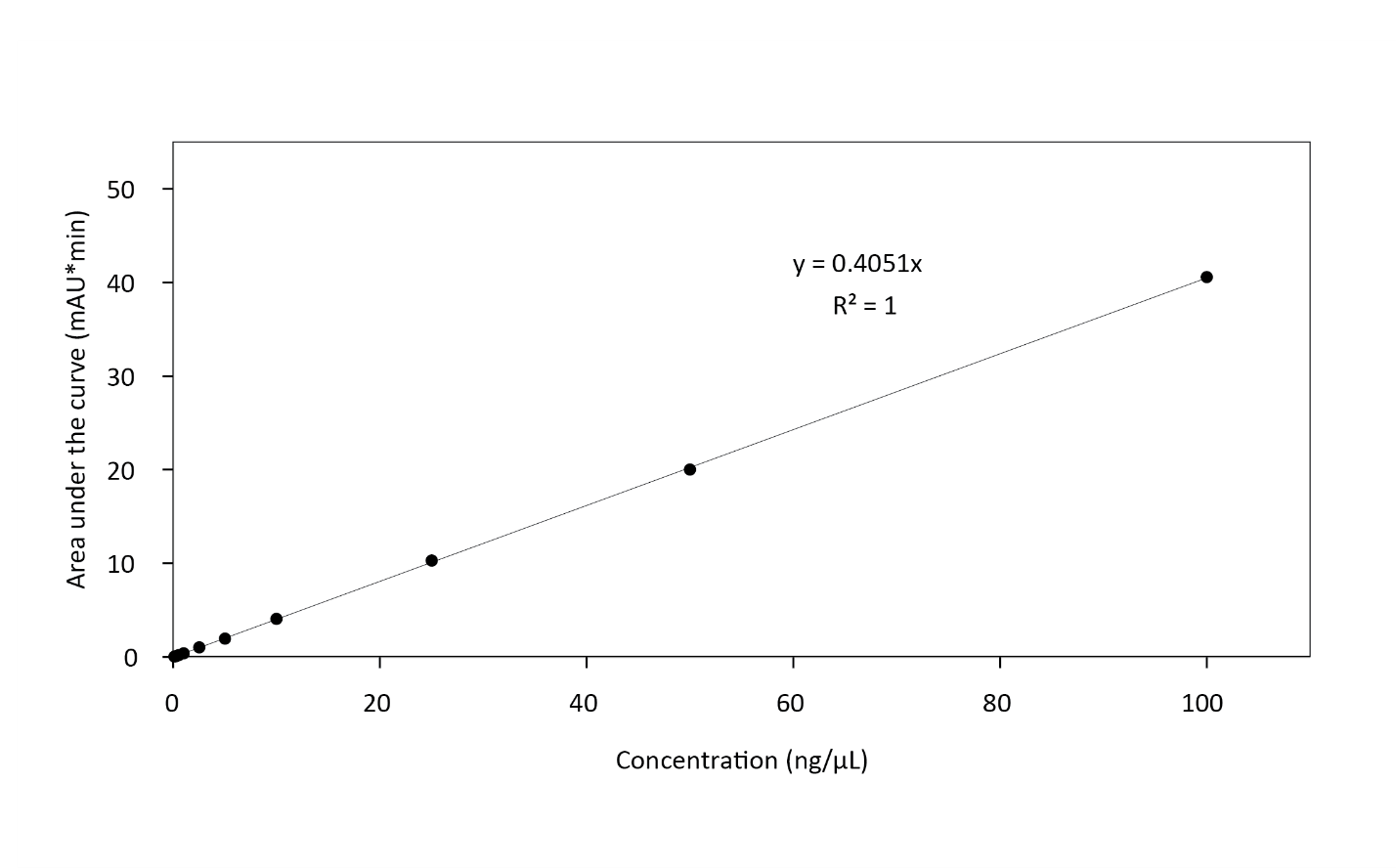

**Fig S5. 4-amino-2,6-dinitrotoluene calibration curve with the Eclipse column.**

**Supplementary Table S1: *D. melanogaster* strains used in this study**

| **Strain** | **Description** | **Reference** | **BDSC#** |
| --- | --- | --- | --- |
| Chr2 AttP3 for microinjections | y1 w1118; PBac{y[+]‐attP‐3B}VK00037 | ^1^ | 9752 |
| Chr3 AttP3 for microinjections | y1 w1118; PBac{y[+]‐attP‐3B}VK00037 | ^1^ | 9748 |
| w^1118^ | w[1118] | ^2^ | 5905 |
| Fly/NsfI | y^1^ w^1118^; PBac{NsfI}attP-3B | This study | N/A |
| Fly/NsfA | y^1^ w^1118;^ PBac{NsfA}attP-3B | This study | N/A |
| Fly/PnrA | y^1^ w^1118;^ PBac{PnrA}attP-3B | This study | N/A |
| Fly/SnrA | y^1^ w^1118;^ PBac{SnrA}attP-3B | This study | N/A |
| Fly/Mm-nr | y^1^ w^1118;^ PBac{Mm-nr}attP-3B | This study | N/A |
| Fly/XenA | y^1^ w^1118;^ PBac{XenA}attP-3B | This study | N/A |

**Supplementary Table S2: primers used in this study**

| Name | **Sequence (5’** 🡪 **3’)** | **Description** |
| --- | --- | --- |
| 2ndconstruct_F_2.0 | cgtttgtcaagcctcatagc | pKT111.[ ] Sanger sequencing |
| 2ndconstruct_R_2.0 | ttcattttatgtttcaggttcaggg | pKT111.[ ] Sanger sequencing |
| DPE5_PnrA_rtPCR_F | ctcgtctggttggttgactggtc | PnrA RT-qPCR assay |
| DPE6_PnrA_rtPCR_R | ctcccactgtatacgactccagatagtc | PnrA RT-qPCR assay |
| DPE7_Mm-NR_rtPCR_F | gagcaagaagataaagatggaaggtacactacc | MmNR RT-qPCR assay |
| DPE8_Mm-NR_rtPCR_R | ccagagtttccttgtgcaaattgacaaaaaag | MmNR RT-qPCR assay |
| DPE9_NsfA_rtPCR_F | caattgctcttgggtgtagtggacac | NsfA RT-qPCR assay |
| DPE10_NsfA_rtPCR_R | gcaggcctcctatataaacacctccc | NsfA RT-qPCR assay |
| DPE13_NsfI_rtPCR_F | gaccaagaggaggcggatgga | PnrA RT-qPCR assay |
| DPE14_NsfI_rtPCR_R | tcaagtccaccctatgcatgtcagc | PnrA RT-qPCR assay |
| DPE15_SnrA_rtPCR_F | cagttgctcttgggagtcgtgg | SnrA RT-qPCR assay |
| DPE16_SnrA_rtPCR_R | ggatgccaccgatatatacgcctcc | SnrA RT-qPCR assay |
| DPE17_XenA_rtPCR_F | attgcgcatgcgggacgtaa | XenA RT-qPCR assay |
| DPE18_XenA_rtPCR_R | atggggcgattgtttcccagc | XenA RT-qPCR assay |

**Supplementary Table S3: plasmids used in this study**

| **Plasmid** | **Integrated into fly strain** | **Description** | **Reference** |
| --- | --- | --- | --- |
| pMC-1-1-1 | N/A | Empty vector, with short alpha-tubulin promoter and SV40 regulatory sequences | ^3^ |
| pKT111.NsfI | Fly/NsfI | Short alpha-tubulin promoter driving  expression of *nsfI* from *E. cloacae* | This study |
| pKT111.NsfA | Fly/NsfA | Short alpha-tubulin promoter driving  expression of *nsfA* from *E. coli* | This study |
| pKT111.PnrA | Fly/PnrA | Short alpha-tubulin promoter driving  expression of *pnrA* from *P. putida* | This study |
| pKT111.SnrA | Fly/SnrA | Short alpha-tubulin promoter driving  expression of *snrA* from *S. enterica* | This study |
| pKT111.Mm-nr | Fly/Mm-nr | Short alpha-tubulin promoter driving  expression of *Mm-nr* from *M. morganii* | This study |
| pKT111.XenA | Fly/XenA | Short alpha-tubulin promoter driving  expression of *xenA* from *P. putida* II-B | This study |

**Supplementary Table S4: plasmid DNA sequences**

| **Plasmid name** | **Sequence (circularized)** |
| --- | --- |
| pKT111.NsfI | AAAATAATCAATTAGGGGTTCCGCGCAACAATTTCCCCGAAAAAAGTGCCCACCTAAATTGTAAAGCGTTTAATATTTTTGTTTAAAATTCGCGTTAAATTTTTTGTTAAATCAGCTCATTTTTTTAACCAATAGGCCGAAAATCGGCAAAAATCCCTTATAAATCAAAAAGAATAGACCGAGATAGGGTTGAGTGTTGTTCCAGTTTGGAACAAGAGTCCACTATTAAAGAACGTGGACTCCAACGTCAAAGGGCGAAAAACCGTCTATCAGGGCGATGGCCCACTACGTGAACCATCACCCTAATCAAGTTTTTTGGGGTCGAGGTGCCGTAAAGCACTAAATCGGAACCCTAAAGGGAGCCCCCGATTTAGAGCTTGACGGGGAAAGCCGGCGAACGTGGCGAGAAAGGAAGGGAAGAAAGCGAAAGGAGCGGGCGCTAGGGCGCTGGCAAGTGTAGCGGTCACGCTGCGCGTAACCACCACACCCGCCGCGCTTAATGCGCCGCTACAGGGCGCGTCCCATTCGCCATTCAGGCTGCGCAACTGTTGGGAAGGGCGATCGGTGCGGGCCTCTTCGCTATTACGCCAGCTGGCGAAAGGGGGATGTGCTGCAAGGCGATTAAGTTGGGTAACGCCAGGGTTTTCCCAGTCACGACGTTGTAAAACGACGGCCAGTGAATTGTAATACGACTCACTATAGGGCGAATTGGGTACGTACCGGGCCCCTAGTATGTATGTAAGTTAATAAAACCCATTTTTGCGGAAAGTAGATAAAAAAAACATTTTTTTTTTTTACTGCACTGGATATCATTGAACTTATCTGATCAGTTTTAAATTTACTTCGATCCAAGGGTATTTGATGTACCAGGTTCTTTCGATTACCTCTCACTCAAAATGACATTCCACTCAAAGTCAGCGCTGTTTGCCTCCTTCTCTGTCCACAGAAATATCGCCGTCTCTTTCGCCGCTGCGTCCGCTATCTCTTTCGCCACCGTTTGTAGCGTTACGTAGCGTCAATGTCCGCCTTCAGTTGCATTTTGTCAGCGGTTTCGTGACGAAGCTCCAAGCGGTTTACGCCATCAATTAAACACAAAGTGCTGTGCCAAAACTCCTCTCGCTTCTTATTTTTGTTTGTTTTTTGAGTGATTGGGGTGGTGATTGGTTTTGGGTGGGTAAGCAGGGGAAAGTGTGAAAAATCCCGGCAATGGGCCAAGAGGATCAGGAGCTATTAATTCGCGGAGGCAGCAAACACCCATCTGCCGAGCATCTGAACAATGTGAGTAGTACATGTGCATACATCTTAAGTTCACTTGATCTATAGGAACTGCGATTGCAACATCAAATTGTCTGCGGCGTGAGAACTGCGACCCACAAAAATCCCAAACCGCAATTGCACAAACAAATAGTGACACGAAACAGATTATTCTGGTAGCTGTTCTCGCTATATAAGACAATTTTTGAGATCATATCATGATCAAGACATCTAAAGGCATTCATTTTCGACTATATTCTTTTTTACAAAAAATATAACAACCAGATATTTTAAGCTGATCCTAGATGCACAAAAAATAAATAAAAGTATAAACCTACTTCGTAGGATACTTCGGGGTACTTTTTGTTCGGGGTTAGATGAGCATAACGCTTGTAGTTGATATTTGAGATCCCCTATCATTGCAGGGTGACAGCGGAGCGGCTTCGCAGAGCTGCATTAACCAGGGCTTCGGGCAGGCCAAAAACTACGGCACGCTCCGGCCACCCAGTCCGCCGGAGGACTCCGGTTCAGGGAGCGGCCAACTAGCCGAGAACCTCACCTATGCCTGGCACAATATGGACATCTTTGGGGCGGTCAATCAGCCGGGCTCCGGATGGCGGCAGCTGGTCAACCGGACACGCGGACTATTCTGCAACGAGCGACACATACCGGCGCCCAGGAAACATTTGCTCAAGAACGGTGAGTTTCTATTCGCAGTCGGCTGATCTGTGTGAAATCTTAATAAAGGGTCCAATTACCAATTTGAAACTCAGTTTGCGGCGTGGCCTATCCGGGCGAACTTTTGGCCGTGATGGGCAGTTCCGGTGCCGGAAAGACGACCCTGCTGAATGCCCTTGCCTTTCGATCGCCGCAGGGCATCCAAGTATCGCCATCCGGGATGCGACTGCTCAATGGCCAACCTGTGGACGCCAAGGAGATGCAGGCCAGGTGCGCCTATGTCCAGCAGGATGACCTCTTTATCGGCTCCCTAACGGCCAGGGAACACCTGATTTTCCAAGCCATGGTGCGGATGCCACGACATCTGACCTATCGGCAGCGAGTGGCCCGCGTGGATCAGGTGATCCAGGAGCTTTCGCTCAGCAAATGTCAGCACACGATCATCGGTGTGCCCGGCAGGGTGAAAGGTCTGTCCGGCGGAGAAAGGAAGCGTCTGGCATTCGCCTCCGAGGCTCTAACCGATCCGCCGCTTCTGATCTGCGATGAGCCCACCTCCGGACTGGACTCCTTTACCGCCCACAGCGTCGTCCAGGTGCTGAAGAAGCTGTCGCAGAAGGGCAAGACCGTCATCCTGACCATTCATCAGCCGTCTTCCGAGCTGTTTGAGCTCTTTGACAAGATCCTTCTGATGGCCGAGGGCAGGGTAGCTTTCTTGGGCACTCCCAGCGAAGCCGTCGACTTCTTTTCCTAGTGAGTTCGATGTGTTTATTAAGGGTATCTAGTATTACATAACATCTCAACTCCTATCCAGCGTGGGTGCCCAGTGTCCTACCAACTACAATCCGGCGGACTTTTACGTACAGGTGTTGGCCGTTGTGCCCGGACGGGAGATCGAGTCCCGTGATCGGATCGCCAAGATATGCGACAATTTTGCCATTAGCAAAGTAGCCCGGGATATGGAGCAGTTGTTGGCCACCAAAAATCTGGAGAAGCCACTGGAGCAGCCGGAGAATGGGTACACCTACAAGGCCACCTGGTTCATGCAGTTCCGGGCGGTCCTGTGGCGATCCTGGCTGTCGGTGCTCAAGGAACCACTCCTCGTAAAAGTGCGACTTATTCAGACAACGGTGAGTGGTTCCAGTGGAAACAAATGATATAACGCTTACAATTCTTGGAAACAAATTCGCTAGATTTTAGATAGAATTGCCTGATTCCACACCCTTCTTAGTTTTTTTCAATGAGATGTATAGTTTATAGTTTTGCAGAAGATAAATAAATTTCATTTAACTCGCGAATATTAATGAGATGCGAGTAACATTTTAATTTGCAGATGGTTGCCATCTTGATTGGCCTCATCTTTTTGGGCCAACAACTCACGCAAGTGGGTGTGATGAATATCAACGGAGCCATCTTCCTCTTCCTGACCAACATGACCTTTCAAAACGTCTTTGCCACGATAAATGTAAGTCATGTTTAGAATACATTTGCATTTCAATAATTTACTAACTTTCTAATGAATCGATTCGATTTAGGTGTTCACCTCAGAGCTGCCAGTTTTTATGAGGGAGGCCCGAAGTCGACTTTATCGCTGTGACACATACTTTCTGGGCAAAACGATTGCCGAATTGCCGCTTTTTCTCACAGTGCCACTGGTCTTCACGGCGATTGCCTATCCGATGATCGGACTGCGGGCCGGAGTGCTGCACTTCTTCAACTGCCTGGCGCTGGTCACTCTGGTGGCCAATGTGTCAACGTCCTTCGGATATCTAATATCCTGCGCCAGCTCCTCGACCTCGATGGCGCTGTCTGTGGGTCCGCCGGTTATCATACCATTCCTGCTCTTTGGCGGCTTCTTCTTGAACTCGGGCTCGGTGCCAGTATACCTCAAATGGTTGTCGTACCTCTCATGGTTCCGTTACGCCAACGAGGGTCTGCTGATTAACCAATGGGCGGACGTGGAGCCGGGCGAAATTAGCTGCACATCGTCGAACACCACGTGCCCCAGTTCGGGCAAGGTCATCCTGGAGACGCTTAACTTCTCCGCCGCCGATCTGCCGCTGGACTACGTGGGTCTGGCCATTCTCATCGTGAGCTTCCGGGTGCTCGCATATCTGGCTCTAAGACTTCGGGCCCGACGCAAGGAGTAGCCGACATATATCCGAAATAACTGCTTGTTTTTTTTTTTTACCATTATTACCATCGTGTTTACTGTTTATTGCCCCCTCAAAAAGCTAATGTAATTATATTTGTGCCAATAAAAACAAGATATGACCTATAGAATACAAGTATTTCCCCTTCGAACATCCCCACAAGTAGACTTTGGATTTGTCTTCTAACCAAAAGACTTACACACCTGCATACCTTACATCAAAAACTCGTTTATCGCTACATAAAACACCGGGATATATTTTTTATATACATACTTTTCAAATCGCGCGCCCTCTTCATAATTCACCTCCACCACACCACGTTTCGTAGTTGCTCTTTCGCTGTCTCCCACCCGCTCTCCGCAACACATTCACCTTTTGTTCGACGACCTTGGAGCGACTGTCGTTAGTTCCGCGCGATTCGGTTCGCTCAAATGGTTCCGAGTGGTTCATTTCGTCTCAATAGAAATTAGTAATAAATATTTGTATGTACAATTTATTTGCTCCAATATATTTGTATATATTTCCCTCACAGCTATATTTATTCTAATTTAATATTATGACTTTTTAAGGTAATTTTTTGTGACCTGTTCGGAGTGATTAGCGTTACAATTTGAACTGAAAGTGACATCCAGTGTTTGTTCCTTGTGTAGATGCATCTCAAAAAAATGGTGGGCATAATAGTGTTGTTTATATATATCAAAAATAACAACTATAATAATAAGAATACATTTAATTTAGAAAATGCTTGGATTTCACTGGAACTAGGCTAGCATAACTTCGTATAATGTATGCTATACGAAGTTATGCTAGCGGATCCGGGAATTGGGAATTCACGTAAGTACTGTCTGCAGCGTAAGCTTCGTACGTAGCGACCGTCTCAAAGTACTGCCTTTCTGCGTTGGAAAACATCGCCTTTTTCGTCCAAAAGGAGTCCCCAGGTTCGATCCGCATGGCGTTGTGCGTGCGTGCCTTTCTTTTCAAATGATTACGGCTATTAACTTGGGGGCGTTAAGTTGGAAACACGTAAATTGCAGACTGCGATTAGAGTGACCATGAGTAGGAGTTCAAAATCTCCTGACATCATTTTCTTAAAACCTGCTTTGTTTTTTACATTTCTATTTAATATAACTCCTATTTGAATAAAAAAACAAAACAAGTTTAGATGTTAAGATATTAACTACATCCTTTGCTCCAAAGGGAGAGGGGAAGTTATGGAGTTAATTAATTTGCTGTTGGAAATCAATATGGAGTCAGAAATATAATGATTTACTAAACCTTATTGAATCGGTAACGATGCGAATTTATATTAAAATAGCTTTTATGAAACATTCAACAAAAATATTATTAATGTTGGCCCACTTTAGCAACCGGTTAGGTCTACCGGTTGGGCAAGCAAAGATTCACGCCCTGGTTCGAGTCCCAACTAGTCCTGCAAAATACCGCAGCAAGTTTTAGAGAGACCAAGTGCCATTACCTCTCCCACTTCAGTTATCGGTTATGCGGCGTTTAAGTCGACAGCTTGCCGTCTCTAGCTCCGGTGCCTATATAAAGCAGCCCGCTTTCCACATTTCATATTCGTTTTACGTTTGTCAAGCCTCATAGCCGGCAGTTCGAACGTATACGCTCTCTGAGTCAGACCTCGAAATCGTAGCTCTACACAATTCTGTGAATTTTCCTTGTCGCGTGTGAAACACTTCCAAGGCAATCTTACAAAATGGACATCATTAGCGTAGCCTTGAAAAGGCATTCCACCAAAGCATTCGATGCGTCCAAAAAGCTCACTGCAGAAGAGGCAGAAAAGATCAAAACACTCCTGCAGTACTCCCCATCCTCCACCAATAGCCAGCCATGGCACTTCATTGTCGCCTCGACTGAAGAAGGTAAAGCTAGGGTAGCAAAGAGCGCGGCAGGTACATACGTCTTTAACGAACGTAAGATGTTGGACGCGAGCCATGTAGTCGTCTTTTGTGCTAAAACTGCCATGGATGATGCGTGGTTGGAACGAGTCGTGGACCAAGAGGAGGCGGATGGACGATTTAACACACCTGAGGCCAAGGCCGCGAATCACAAGGGACGCACATACTTTGCTGACATGCATAGGGTGGACTTGAAGGACGATGATCAGTGGATGGCAAAACAGGTATATTTGAACGTAGGCAACTTTCTCCTGGGCGTTGGAGCAATGGGATTGGACGCGGTTCCTATCGAAGGATTTGACGCTGCGATTCTGGATGAAGAATTTGGTTTGAAGGAGAAGGGTTTTACTTCCCTCGTCGTAGTGCCCGTCGGTCACCATTCCGTGGAAGACTTTAACGCGACGCTCCCGAAGTCGAGGTTGCCTCTCTCCACAATTGTCACGGAATGCTAAGCCCGACTCTAGATCATAATCAGCCATACCACATTTGTAGAGGTTTTACTTGCTTTAAAAAACCTCCCACACCTCCCCCTGAACCTGAAACATAAAATGAATGCAATTGTTGTTGTTAACTTGTTTATTGCAGCTTATAATGGTTACAAATAAAGCAATAGCATCACAAATTTCACAAATAAAGCATTTTTTTCACTGCATTCTAGTTGTGGTTTGCCCAAACTCATCAATGTATCTTAGCGGCTCGAGGGTACCTCTAGAGATCCACTAGTGTCGACGATGTAGGTCACGGTCTCGAAGCCGCGGTGCGGGTGCCAGGGCGTGCCCTTGGGCTCCCCGGGCGCGTACTCCACCTCACCCATCTGGTCCATCATGATGAACGGGTCGAGGTGGCGGTAGTTGATCCCGGCGAACGCGCGGCGCACCGGGAAGCCCTCGCCCTCGAAACCGCTGGGCGCGGTGGTCACGGTGAGCACGGGACGTGCGACGGCGTCGGCGGGTGCGGATACGCGGGGCAGCGTCAGCGGGTTCTCGACGGTCACGGCGGGCATGTCGACACTAGTTCTAGCCAGCTTTTGTTCCCTTTAGTGAGGGTTAATTTCGAGCTTGGCGTAATCATGGTCATAGCTGTTTCCTGTGTGAAATTGTTATCCGCTCACAATTCCACACAACATACGAGCCGGAAGCATAAAGTGTAAAGCCTGGGGTGCCTAATGAGTGAGCTAACTCACATTAATTGCGTTGCGCTCACTGCCCGCTTTCCAGTCGGGAAACCTGTCGTGCCAGCTGCATTAATGAATCGGCCAACGCGCGGGGAGAGGCGGTTTGCGTATTGGGCGCTCTTCCGCTTCCTCGCTCACTGACTCGCTGCGCTCGGTCGTTCGGCTGCGGCGAGCGGTATCAGCTCACTCAAAGGCGGTAATACGGTTATCCACAGAATCAGGGGATAACGCAGGAAAGAACATGTGAGCAAAAGGCCAGCAAAAGGCCAGGAACCGTAAAAAGGCCGCGTTGCTGGCGTTTTTCCATAGGCTCCGCCCCCCTGACGAGCATCACAAAAATCGACGCTCAAGTCAGAGGTGGCGAAACCCGACAGGACTATAAAGATACCAGGCGTTTCCCCCTGGAAGCTCCCTCGTGCGCTCTCCTGTTCCGACCCTGCCGCTTACCGGATACCTGTCCGCCTTTCTCCCTTCGGGAAGCGTGGCGCTTTCTCATAGCTCACGCTGTAGGTATCTCAGTTCGGTGTAGGTCGTTCGCTCCAAGCTGGGCTGTGTGCACGAACCCCCCGTTCAGCCCGACCGCTGCGCCTTATCCGGTAACTATCGTCTTGAGTCCAACCCGGTAAGACACGACTTATCGCCACTGGCAGCAGCCACTGGTAACAGGATTAGCAGAGCGAGGTATGTAGGCGGTGCTACAGAGTTCTTGAAGTGGTGGCCTAACTACGGCTACACTAGAAGAACAGTATTTGGTATCTGCGCTCTGCTGAAGCCAGTTACCTTCGGAAAAAGAGTTGGTAGCTCTTGATCCGGCAAACAAACCACCGCTGGTAGCGGTGGTTTTTTTGTTTGCAAGCAGCAGATTACGCGCAGAAAAAAAGGATCTCAAGAAGATCCTTTGATCTTTTCTACGGGGTCTGACGCTCAGTGGAACGAAAACTCACGTTAAGGGATTTTGGTCATGAGATTATCAAAAAGGATCTTCACCTAGATCCTTTTAAATTAAAAATGAAGTTTTAAATCAATCTAAAGTATATATGAGTAAACTTGGTCTGACAGTTACCAATGCTTAATCAGTGAGGCACCTATCTCAGCGATCTGTCTATTTCGTTCATCCATAGTTGCCTGACTCCCCGTCGTGTAGATAACTACGATACGGGAGGGCTTACCATCTGGCCCCAGTGCTGCAATGATACCGCGAGACCCACGCTCACCGGCTCCAGATTTATCAGCAATAAACCAGCCAGCCGGAAGGGCCGAGCGCAGAAGTGGTCCTGCAACTTTATCCGCCTCCATCCAGTCTATTAATTGTTGCCGGGAAGCTAGAGTAAGTAGTTCGCCAGTTAATAGTTTGCGCAACGTTGTTGCCATTGCTACAGGCATCGTGGTGTCACGCTCGTCGTTTGGTATGGCTTCATTCAGCTCCGGTTCCCAACGATCAAGGCGAGTTACATGATCCCCCATGTTGTGCAAAAAAGCGGTTAGCTCCTTCGGTCCTCCGATCGTTGTCAGAAGTAAGTTGGCCGCAGTGTTATCACTCATGGTTATGGCAGCACTGCATAATTCTCTTACTGTCATGCCATCCGTAAGATGCTTTTCTGTGACTGGTGAGTACTCAACCAAGTCATTCTGAGAATAGTGTATGCGGCGACCGAGTTGCTCTTGCCCGGCGTCAATACGGGATAATACCGCGCCACATAGCAGAACTTTAAAAGTGCTCATCATTGGAAAACGTTCTTCGGGGCGAAAACTCTCAAGGATCTTACCGCTGTTGAGATCCAGTTCGATGTAACCCACTCGTGCACCCAACTGATCTTCAGCATCTTTTACTTTCACCAGCGTTTCTGGGTGAGCAAAAACAGGAAGGCAAAATGCCGCAAAAAAGGGAATAAGGGCGACACGGAAATGTTGAATACTCATACTCTTCCTTTTTCAATATTATTGAAGCATTTATCAGGGTTATTGTCTCATGAGCGGATACATATTTGAATGTATTTAGAAAAATAAACAAATAGGGGTTCCGCGCACATTTCCTCCGAAAAGTGCCACCTAAATTGTAAGCGTTAATATTTTTGTTAAAATTCGCGTTAAATTTTTTGTTAAATCAGCTCATTTTTTTAACCAATAGGCCGAAATCGGCAAAAATCCCCTTATAAATCAAAAAGAATAGACCGAGATAGGGTTGAGTGTTGTTCCCAGTTTTGGC |
| pKT111.NsfA | AAAATAATCAATTAGGGGTTCCGCGCAACAATTTCCCCGAAAAAAGTGCCCACCTAAATTGTAAAGCGTTTAATATTTTTGTTTAAAATTCGCGTTAAATTTTTTGTTAAATCAGCTCATTTTTTTAACCAATAGGCCGAAAATCGGCAAAAATCCCTTATAAATCAAAAAGAATAGACCGAGATAGGGTTGAGTGTTGTTCCAGTTTGGAACAAGAGTCCACTATTAAAGAACGTGGACTCCAACGTCAAAGGGCGAAAAACCGTCTATCAGGGCGATGGCCCACTACGTGAACCATCACCCTAATCAAGTTTTTTGGGGTCGAGGTGCCGTAAAGCACTAAATCGGAACCCTAAAGGGAGCCCCCGATTTAGAGCTTGACGGGGAAAGCCGGCGAACGTGGCGAGAAAGGAAGGGAAGAAAGCGAAAGGAGCGGGCGCTAGGGCGCTGGCAAGTGTAGCGGTCACGCTGCGCGTAACCACCACACCCGCCGCGCTTAATGCGCCGCTACAGGGCGCGTCCCATTCGCCATTCAGGCTGCGCAACTGTTGGGAAGGGCGATCGGTGCGGGCCTCTTCGCTATTACGCCAGCTGGCGAAAGGGGGATGTGCTGCAAGGCGATTAAGTTGGGTAACGCCAGGGTTTTCCCAGTCACGACGTTGTAAAACGACGGCCAGTGAATTGTAATACGACTCACTATAGGGCGAATTGGGTACGTACCGGGCCCCTAGTATGTATGTAAGTTAATAAAACCCATTTTTGCGGAAAGTAGATAAAAAAAACATTTTTTTTTTTTACTGCACTGGATATCATTGAACTTATCTGATCAGTTTTAAATTTACTTCGATCCAAGGGTATTTGATGTACCAGGTTCTTTCGATTACCTCTCACTCAAAATGACATTCCACTCAAAGTCAGCGCTGTTTGCCTCCTTCTCTGTCCACAGAAATATCGCCGTCTCTTTCGCCGCTGCGTCCGCTATCTCTTTCGCCACCGTTTGTAGCGTTACGTAGCGTCAATGTCCGCCTTCAGTTGCATTTTGTCAGCGGTTTCGTGACGAAGCTCCAAGCGGTTTACGCCATCAATTAAACACAAAGTGCTGTGCCAAAACTCCTCTCGCTTCTTATTTTTGTTTGTTTTTTGAGTGATTGGGGTGGTGATTGGTTTTGGGTGGGTAAGCAGGGGAAAGTGTGAAAAATCCCGGCAATGGGCCAAGAGGATCAGGAGCTATTAATTCGCGGAGGCAGCAAACACCCATCTGCCGAGCATCTGAACAATGTGAGTAGTACATGTGCATACATCTTAAGTTCACTTGATCTATAGGAACTGCGATTGCAACATCAAATTGTCTGCGGCGTGAGAACTGCGACCCACAAAAATCCCAAACCGCAATTGCACAAACAAATAGTGACACGAAACAGATTATTCTGGTAGCTGTTCTCGCTATATAAGACAATTTTTGAGATCATATCATGATCAAGACATCTAAAGGCATTCATTTTCGACTATATTCTTTTTTACAAAAAATATAACAACCAGATATTTTAAGCTGATCCTAGATGCACAAAAAATAAATAAAAGTATAAACCTACTTCGTAGGATACTTCGGGGTACTTTTTGTTCGGGGTTAGATGAGCATAACGCTTGTAGTTGATATTTGAGATCCCCTATCATTGCAGGGTGACAGCGGAGCGGCTTCGCAGAGCTGCATTAACCAGGGCTTCGGGCAGGCCAAAAACTACGGCACGCTCCGGCCACCCAGTCCGCCGGAGGACTCCGGTTCAGGGAGCGGCCAACTAGCCGAGAACCTCACCTATGCCTGGCACAATATGGACATCTTTGGGGCGGTCAATCAGCCGGGCTCCGGATGGCGGCAGCTGGTCAACCGGACACGCGGACTATTCTGCAACGAGCGACACATACCGGCGCCCAGGAAACATTTGCTCAAGAACGGTGAGTTTCTATTCGCAGTCGGCTGATCTGTGTGAAATCTTAATAAAGGGTCCAATTACCAATTTGAAACTCAGTTTGCGGCGTGGCCTATCCGGGCGAACTTTTGGCCGTGATGGGCAGTTCCGGTGCCGGAAAGACGACCCTGCTGAATGCCCTTGCCTTTCGATCGCCGCAGGGCATCCAAGTATCGCCATCCGGGATGCGACTGCTCAATGGCCAACCTGTGGACGCCAAGGAGATGCAGGCCAGGTGCGCCTATGTCCAGCAGGATGACCTCTTTATCGGCTCCCTAACGGCCAGGGAACACCTGATTTTCCAAGCCATGGTGCGGATGCCACGACATCTGACCTATCGGCAGCGAGTGGCCCGCGTGGATCAGGTGATCCAGGAGCTTTCGCTCAGCAAATGTCAGCACACGATCATCGGTGTGCCCGGCAGGGTGAAAGGTCTGTCCGGCGGAGAAAGGAAGCGTCTGGCATTCGCCTCCGAGGCTCTAACCGATCCGCCGCTTCTGATCTGCGATGAGCCCACCTCCGGACTGGACTCCTTTACCGCCCACAGCGTCGTCCAGGTGCTGAAGAAGCTGTCGCAGAAGGGCAAGACCGTCATCCTGACCATTCATCAGCCGTCTTCCGAGCTGTTTGAGCTCTTTGACAAGATCCTTCTGATGGCCGAGGGCAGGGTAGCTTTCTTGGGCACTCCCAGCGAAGCCGTCGACTTCTTTTCCTAGTGAGTTCGATGTGTTTATTAAGGGTATCTAGTATTACATAACATCTCAACTCCTATCCAGCGTGGGTGCCCAGTGTCCTACCAACTACAATCCGGCGGACTTTTACGTACAGGTGTTGGCCGTTGTGCCCGGACGGGAGATCGAGTCCCGTGATCGGATCGCCAAGATATGCGACAATTTTGCCATTAGCAAAGTAGCCCGGGATATGGAGCAGTTGTTGGCCACCAAAAATCTGGAGAAGCCACTGGAGCAGCCGGAGAATGGGTACACCTACAAGGCCACCTGGTTCATGCAGTTCCGGGCGGTCCTGTGGCGATCCTGGCTGTCGGTGCTCAAGGAACCACTCCTCGTAAAAGTGCGACTTATTCAGACAACGGTGAGTGGTTCCAGTGGAAACAAATGATATAACGCTTACAATTCTTGGAAACAAATTCGCTAGATTTTAGATAGAATTGCCTGATTCCACACCCTTCTTAGTTTTTTTCAATGAGATGTATAGTTTATAGTTTTGCAGAAGATAAATAAATTTCATTTAACTCGCGAATATTAATGAGATGCGAGTAACATTTTAATTTGCAGATGGTTGCCATCTTGATTGGCCTCATCTTTTTGGGCCAACAACTCACGCAAGTGGGTGTGATGAATATCAACGGAGCCATCTTCCTCTTCCTGACCAACATGACCTTTCAAAACGTCTTTGCCACGATAAATGTAAGTCATGTTTAGAATACATTTGCATTTCAATAATTTACTAACTTTCTAATGAATCGATTCGATTTAGGTGTTCACCTCAGAGCTGCCAGTTTTTATGAGGGAGGCCCGAAGTCGACTTTATCGCTGTGACACATACTTTCTGGGCAAAACGATTGCCGAATTGCCGCTTTTTCTCACAGTGCCACTGGTCTTCACGGCGATTGCCTATCCGATGATCGGACTGCGGGCCGGAGTGCTGCACTTCTTCAACTGCCTGGCGCTGGTCACTCTGGTGGCCAATGTGTCAACGTCCTTCGGATATCTAATATCCTGCGCCAGCTCCTCGACCTCGATGGCGCTGTCTGTGGGTCCGCCGGTTATCATACCATTCCTGCTCTTTGGCGGCTTCTTCTTGAACTCGGGCTCGGTGCCAGTATACCTCAAATGGTTGTCGTACCTCTCATGGTTCCGTTACGCCAACGAGGGTCTGCTGATTAACCAATGGGCGGACGTGGAGCCGGGCGAAATTAGCTGCACATCGTCGAACACCACGTGCCCCAGTTCGGGCAAGGTCATCCTGGAGACGCTTAACTTCTCCGCCGCCGATCTGCCGCTGGACTACGTGGGTCTGGCCATTCTCATCGTGAGCTTCCGGGTGCTCGCATATCTGGCTCTAAGACTTCGGGCCCGACGCAAGGAGTAGCCGACATATATCCGAAATAACTGCTTGTTTTTTTTTTTTACCATTATTACCATCGTGTTTACTGTTTATTGCCCCCTCAAAAAGCTAATGTAATTATATTTGTGCCAATAAAAACAAGATATGACCTATAGAATACAAGTATTTCCCCTTCGAACATCCCCACAAGTAGACTTTGGATTTGTCTTCTAACCAAAAGACTTACACACCTGCATACCTTACATCAAAAACTCGTTTATCGCTACATAAAACACCGGGATATATTTTTTATATACATACTTTTCAAATCGCGCGCCCTCTTCATAATTCACCTCCACCACACCACGTTTCGTAGTTGCTCTTTCGCTGTCTCCCACCCGCTCTCCGCAACACATTCACCTTTTGTTCGACGACCTTGGAGCGACTGTCGTTAGTTCCGCGCGATTCGGTTCGCTCAAATGGTTCCGAGTGGTTCATTTCGTCTCAATAGAAATTAGTAATAAATATTTGTATGTACAATTTATTTGCTCCAATATATTTGTATATATTTCCCTCACAGCTATATTTATTCTAATTTAATATTATGACTTTTTAAGGTAATTTTTTGTGACCTGTTCGGAGTGATTAGCGTTACAATTTGAACTGAAAGTGACATCCAGTGTTTGTTCCTTGTGTAGATGCATCTCAAAAAAATGGTGGGCATAATAGTGTTGTTTATATATATCAAAAATAACAACTATAATAATAAGAATACATTTAATTTAGAAAATGCTTGGATTTCACTGGAACTAGGCTAGCATAACTTCGTATAATGTATGCTATACGAAGTTATGCTAGCGGATCCGGGAATTGGGAATTCACGTAAGTACTGTCTGCAGCGTAAGCTTCGTACGTAGCGACCGTCTCAAAGTACTGCCTTTCTGCGTTGGAAAACATCGCCTTTTTCGTCCAAAAGGAGTCCCCAGGTTCGATCCGCATGGCGTTGTGCGTGCGTGCCTTTCTTTTCAAATGATTACGGCTATTAACTTGGGGGCGTTAAGTTGGAAACACGTAAATTGCAGACTGCGATTAGAGTGACCATGAGTAGGAGTTCAAAATCTCCTGACATCATTTTCTTAAAACCTGCTTTGTTTTTTACATTTCTATTTAATATAACTCCTATTTGAATAAAAAAACAAAACAAGTTTAGATGTTAAGATATTAACTACATCCTTTGCTCCAAAGGGAGAGGGGAAGTTATGGAGTTAATTAATTTGCTGTTGGAAATCAATATGGAGTCAGAAATATAATGATTTACTAAACCTTATTGAATCGGTAACGATGCGAATTTATATTAAAATAGCTTTTATGAAACATTCAACAAAAATATTATTAATGTTGGCCCACTTTAGCAACCGGTTAGGTCTACCGGTTGGGCAAGCAAAGATTCACGCCCTGGTTCGAGTCCCAACTAGTCCTGCAAAATACCGCAGCAAGTTTTAGAGAGACCAAGTGCCATTACCTCTCCCACTTCAGTTATCGGTTATGCGGCGTTTAAGTCGACAGCTTGCCGTCTCTAGCTCCGGTGCCTATATAAAGCAGCCCGCTTTCCACATTTCATATTCGTTTTACGTTTGTCAAGCCTCATAGCCGGCAGTTCGAACGTATACGCTCTCTGAGTCAGACCTCGAAATCGTAGCTCTACACAATTCTGTGAATTTTCCTTGTCGCGTGTGAAACACTTCCAAGGCAATCTTACAAAATGACCCCTACAATCGAGTTGATTTGCGGTCATCGTTCGATTCGCCATTTTACTGACGAGCCCATATCGGAAGCCCAGCGAGAAGCGATCATAAACAGCGCCCGCGCGACATCCTCCAGCTCGTTTCTCCAATGCTCCAGCATAATCCGTATTACCGATAAAGCCCTGCGAGAAGAACTCGTTACACTCACCGGTGGACAGAAGCATGTTGCTCAGGCTGCTGAGTTTTGGGTATTTTGTGCTGACTTCAACCGACATCTCCAGATCTGTCCCGACGCGCAATTGGGTTTGGCAGAACAATTGCTCTTGGGTGTAGTGGACACTGCGATGATGGCGCAGAATGCTCTGATCGCGGCGGAGTCCCTCGGATTGGGAGGTGTTTATATAGGAGGCCTGCGTAATAACATTGAGGCAGTCACGAAGCTGCTCAAGTTGCCGCAGCATGTCTTGCCTCTCTTCGGATTGTGTCTGGGTTGGCCCGCGGATAATCCCGATCTGAAGCCACGGCTCCCTGCTTCCATCCTGGTCCATGAAAATAGCTACCAACCGTTGGACAAGGGTGCGCTGGCTCAGTATGACGAGCAACTGGCCGAATACTACCTGACTAGGGGATCGAACAATCGCCGAGATACATGGTCGGATCATATTCGGCGAACGATCATTAAAGAGTCCCGGCCGTTTATCTTGGACTACCTGCATAAACAAGGATGGGCAACTCGCTAAGCCCGACTCTAGATCATAATCAGCCATACCACATTTGTAGAGGTTTTACTTGCTTTAAAAAACCTCCCACACCTCCCCCTGAACCTGAAACATAAAATGAATGCAATTGTTGTTGTTAACTTGTTTATTGCAGCTTATAATGGTTACAAATAAAGCAATAGCATCACAAATTTCACAAATAAAGCATTTTTTTCACTGCATTCTAGTTGTGGTTTGCCCAAACTCATCAATGTATCTTAGCGGCTCGAGGGTACCTCTAGAGATCCACTAGTGTCGACGATGTAGGTCACGGTCTCGAAGCCGCGGTGCGGGTGCCAGGGCGTGCCCTTGGGCTCCCCGGGCGCGTACTCCACCTCACCCATCTGGTCCATCATGATGAACGGGTCGAGGTGGCGGTAGTTGATCCCGGCGAACGCGCGGCGCACCGGGAAGCCCTCGCCCTCGAAACCGCTGGGCGCGGTGGTCACGGTGAGCACGGGACGTGCGACGGCGTCGGCGGGTGCGGATACGCGGGGCAGCGTCAGCGGGTTCTCGACGGTCACGGCGGGCATGTCGACACTAGTTCTAGCCAGCTTTTGTTCCCTTTAGTGAGGGTTAATTTCGAGCTTGGCGTAATCATGGTCATAGCTGTTTCCTGTGTGAAATTGTTATCCGCTCACAATTCCACACAACATACGAGCCGGAAGCATAAAGTGTAAAGCCTGGGGTGCCTAATGAGTGAGCTAACTCACATTAATTGCGTTGCGCTCACTGCCCGCTTTCCAGTCGGGAAACCTGTCGTGCCAGCTGCATTAATGAATCGGCCAACGCGCGGGGAGAGGCGGTTTGCGTATTGGGCGCTCTTCCGCTTCCTCGCTCACTGACTCGCTGCGCTCGGTCGTTCGGCTGCGGCGAGCGGTATCAGCTCACTCAAAGGCGGTAATACGGTTATCCACAGAATCAGGGGATAACGCAGGAAAGAACATGTGAGCAAAAGGCCAGCAAAAGGCCAGGAACCGTAAAAAGGCCGCGTTGCTGGCGTTTTTCCATAGGCTCCGCCCCCCTGACGAGCATCACAAAAATCGACGCTCAAGTCAGAGGTGGCGAAACCCGACAGGACTATAAAGATACCAGGCGTTTCCCCCTGGAAGCTCCCTCGTGCGCTCTCCTGTTCCGACCCTGCCGCTTACCGGATACCTGTCCGCCTTTCTCCCTTCGGGAAGCGTGGCGCTTTCTCATAGCTCACGCTGTAGGTATCTCAGTTCGGTGTAGGTCGTTCGCTCCAAGCTGGGCTGTGTGCACGAACCCCCCGTTCAGCCCGACCGCTGCGCCTTATCCGGTAACTATCGTCTTGAGTCCAACCCGGTAAGACACGACTTATCGCCACTGGCAGCAGCCACTGGTAACAGGATTAGCAGAGCGAGGTATGTAGGCGGTGCTACAGAGTTCTTGAAGTGGTGGCCTAACTACGGCTACACTAGAAGAACAGTATTTGGTATCTGCGCTCTGCTGAAGCCAGTTACCTTCGGAAAAAGAGTTGGTAGCTCTTGATCCGGCAAACAAACCACCGCTGGTAGCGGTGGTTTTTTTGTTTGCAAGCAGCAGATTACGCGCAGAAAAAAAGGATCTCAAGAAGATCCTTTGATCTTTTCTACGGGGTCTGACGCTCAGTGGAACGAAAACTCACGTTAAGGGATTTTGGTCATGAGATTATCAAAAAGGATCTTCACCTAGATCCTTTTAAATTAAAAATGAAGTTTTAAATCAATCTAAAGTATATATGAGTAAACTTGGTCTGACAGTTACCAATGCTTAATCAGTGAGGCACCTATCTCAGCGATCTGTCTATTTCGTTCATCCATAGTTGCCTGACTCCCCGTCGTGTAGATAACTACGATACGGGAGGGCTTACCATCTGGCCCCAGTGCTGCAATGATACCGCGAGACCCACGCTCACCGGCTCCAGATTTATCAGCAATAAACCAGCCAGCCGGAAGGGCCGAGCGCAGAAGTGGTCCTGCAACTTTATCCGCCTCCATCCAGTCTATTAATTGTTGCCGGGAAGCTAGAGTAAGTAGTTCGCCAGTTAATAGTTTGCGCAACGTTGTTGCCATTGCTACAGGCATCGTGGTGTCACGCTCGTCGTTTGGTATGGCTTCATTCAGCTCCGGTTCCCAACGATCAAGGCGAGTTACATGATCCCCCATGTTGTGCAAAAAAGCGGTTAGCTCCTTCGGTCCTCCGATCGTTGTCAGAAGTAAGTTGGCCGCAGTGTTATCACTCATGGTTATGGCAGCACTGCATAATTCTCTTACTGTCATGCCATCCGTAAGATGCTTTTCTGTGACTGGTGAGTACTCAACCAAGTCATTCTGAGAATAGTGTATGCGGCGACCGAGTTGCTCTTGCCCGGCGTCAATACGGGATAATACCGCGCCACATAGCAGAACTTTAAAAGTGCTCATCATTGGAAAACGTTCTTCGGGGCGAAAACTCTCAAGGATCTTACCGCTGTTGAGATCCAGTTCGATGTAACCCACTCGTGCACCCAACTGATCTTCAGCATCTTTTACTTTCACCAGCGTTTCTGGGTGAGCAAAAACAGGAAGGCAAAATGCCGCAAAAAAGGGAATAAGGGCGACACGGAAATGTTGAATACTCATACTCTTCCTTTTTCAATATTATTGAAGCATTTATCAGGGTTATTGTCTCATGAGCGGATACATATTTGAATGTATTTAGAAAAATAAACAAATAGGGGTTCCGCGCACATTTCCTCCGAAAAGTGCCACCTAAATTGTAAGCGTTAATATTTTTGTTAAAATTCGCGTTAAATTTTTTGTTAAATCAGCTCATTTTTTTAACCAATAGGCCGAAATCGGCAAAAATCCCCTTATAAATCAAAAAGAATAGACCGAGATAGGGTTGAGTGTTGTTCCCAGTTTTGGC |
| pKT111.PnrA | AAAATAATCAATTAGGGGTTCCGCGCAACAATTTCCCCGAAAAAAGTGCCCACCTAAATTGTAAAGCGTTTAATATTTTTGTTTAAAATTCGCGTTAAATTTTTTGTTAAATCAGCTCATTTTTTTAACCAATAGGCCGAAAATCGGCAAAAATCCCTTATAAATCAAAAAGAATAGACCGAGATAGGGTTGAGTGTTGTTCCAGTTTGGAACAAGAGTCCACTATTAAAGAACGTGGACTCCAACGTCAAAGGGCGAAAAACCGTCTATCAGGGCGATGGCCCACTACGTGAACCATCACCCTAATCAAGTTTTTTGGGGTCGAGGTGCCGTAAAGCACTAAATCGGAACCCTAAAGGGAGCCCCCGATTTAGAGCTTGACGGGGAAAGCCGGCGAACGTGGCGAGAAAGGAAGGGAAGAAAGCGAAAGGAGCGGGCGCTAGGGCGCTGGCAAGTGTAGCGGTCACGCTGCGCGTAACCACCACACCCGCCGCGCTTAATGCGCCGCTACAGGGCGCGTCCCATTCGCCATTCAGGCTGCGCAACTGTTGGGAAGGGCGATCGGTGCGGGCCTCTTCGCTATTACGCCAGCTGGCGAAAGGGGGATGTGCTGCAAGGCGATTAAGTTGGGTAACGCCAGGGTTTTCCCAGTCACGACGTTGTAAAACGACGGCCAGTGAATTGTAATACGACTCACTATAGGGCGAATTGGGTACGTACCGGGCCCCTAGTATGTATGTAAGTTAATAAAACCCATTTTTGCGGAAAGTAGATAAAAAAAACATTTTTTTTTTTTACTGCACTGGATATCATTGAACTTATCTGATCAGTTTTAAATTTACTTCGATCCAAGGGTATTTGATGTACCAGGTTCTTTCGATTACCTCTCACTCAAAATGACATTCCACTCAAAGTCAGCGCTGTTTGCCTCCTTCTCTGTCCACAGAAATATCGCCGTCTCTTTCGCCGCTGCGTCCGCTATCTCTTTCGCCACCGTTTGTAGCGTTACGTAGCGTCAATGTCCGCCTTCAGTTGCATTTTGTCAGCGGTTTCGTGACGAAGCTCCAAGCGGTTTACGCCATCAATTAAACACAAAGTGCTGTGCCAAAACTCCTCTCGCTTCTTATTTTTGTTTGTTTTTTGAGTGATTGGGGTGGTGATTGGTTTTGGGTGGGTAAGCAGGGGAAAGTGTGAAAAATCCCGGCAATGGGCCAAGAGGATCAGGAGCTATTAATTCGCGGAGGCAGCAAACACCCATCTGCCGAGCATCTGAACAATGTGAGTAGTACATGTGCATACATCTTAAGTTCACTTGATCTATAGGAACTGCGATTGCAACATCAAATTGTCTGCGGCGTGAGAACTGCGACCCACAAAAATCCCAAACCGCAATTGCACAAACAAATAGTGACACGAAACAGATTATTCTGGTAGCTGTTCTCGCTATATAAGACAATTTTTGAGATCATATCATGATCAAGACATCTAAAGGCATTCATTTTCGACTATATTCTTTTTTACAAAAAATATAACAACCAGATATTTTAAGCTGATCCTAGATGCACAAAAAATAAATAAAAGTATAAACCTACTTCGTAGGATACTTCGGGGTACTTTTTGTTCGGGGTTAGATGAGCATAACGCTTGTAGTTGATATTTGAGATCCCCTATCATTGCAGGGTGACAGCGGAGCGGCTTCGCAGAGCTGCATTAACCAGGGCTTCGGGCAGGCCAAAAACTACGGCACGCTCCGGCCACCCAGTCCGCCGGAGGACTCCGGTTCAGGGAGCGGCCAACTAGCCGAGAACCTCACCTATGCCTGGCACAATATGGACATCTTTGGGGCGGTCAATCAGCCGGGCTCCGGATGGCGGCAGCTGGTCAACCGGACACGCGGACTATTCTGCAACGAGCGACACATACCGGCGCCCAGGAAACATTTGCTCAAGAACGGTGAGTTTCTATTCGCAGTCGGCTGATCTGTGTGAAATCTTAATAAAGGGTCCAATTACCAATTTGAAACTCAGTTTGCGGCGTGGCCTATCCGGGCGAACTTTTGGCCGTGATGGGCAGTTCCGGTGCCGGAAAGACGACCCTGCTGAATGCCCTTGCCTTTCGATCGCCGCAGGGCATCCAAGTATCGCCATCCGGGATGCGACTGCTCAATGGCCAACCTGTGGACGCCAAGGAGATGCAGGCCAGGTGCGCCTATGTCCAGCAGGATGACCTCTTTATCGGCTCCCTAACGGCCAGGGAACACCTGATTTTCCAAGCCATGGTGCGGATGCCACGACATCTGACCTATCGGCAGCGAGTGGCCCGCGTGGATCAGGTGATCCAGGAGCTTTCGCTCAGCAAATGTCAGCACACGATCATCGGTGTGCCCGGCAGGGTGAAAGGTCTGTCCGGCGGAGAAAGGAAGCGTCTGGCATTCGCCTCCGAGGCTCTAACCGATCCGCCGCTTCTGATCTGCGATGAGCCCACCTCCGGACTGGACTCCTTTACCGCCCACAGCGTCGTCCAGGTGCTGAAGAAGCTGTCGCAGAAGGGCAAGACCGTCATCCTGACCATTCATCAGCCGTCTTCCGAGCTGTTTGAGCTCTTTGACAAGATCCTTCTGATGGCCGAGGGCAGGGTAGCTTTCTTGGGCACTCCCAGCGAAGCCGTCGACTTCTTTTCCTAGTGAGTTCGATGTGTTTATTAAGGGTATCTAGTATTACATAACATCTCAACTCCTATCCAGCGTGGGTGCCCAGTGTCCTACCAACTACAATCCGGCGGACTTTTACGTACAGGTGTTGGCCGTTGTGCCCGGACGGGAGATCGAGTCCCGTGATCGGATCGCCAAGATATGCGACAATTTTGCCATTAGCAAAGTAGCCCGGGATATGGAGCAGTTGTTGGCCACCAAAAATCTGGAGAAGCCACTGGAGCAGCCGGAGAATGGGTACACCTACAAGGCCACCTGGTTCATGCAGTTCCGGGCGGTCCTGTGGCGATCCTGGCTGTCGGTGCTCAAGGAACCACTCCTCGTAAAAGTGCGACTTATTCAGACAACGGTGAGTGGTTCCAGTGGAAACAAATGATATAACGCTTACAATTCTTGGAAACAAATTCGCTAGATTTTAGATAGAATTGCCTGATTCCACACCCTTCTTAGTTTTTTTCAATGAGATGTATAGTTTATAGTTTTGCAGAAGATAAATAAATTTCATTTAACTCGCGAATATTAATGAGATGCGAGTAACATTTTAATTTGCAGATGGTTGCCATCTTGATTGGCCTCATCTTTTTGGGCCAACAACTCACGCAAGTGGGTGTGATGAATATCAACGGAGCCATCTTCCTCTTCCTGACCAACATGACCTTTCAAAACGTCTTTGCCACGATAAATGTAAGTCATGTTTAGAATACATTTGCATTTCAATAATTTACTAACTTTCTAATGAATCGATTCGATTTAGGTGTTCACCTCAGAGCTGCCAGTTTTTATGAGGGAGGCCCGAAGTCGACTTTATCGCTGTGACACATACTTTCTGGGCAAAACGATTGCCGAATTGCCGCTTTTTCTCACAGTGCCACTGGTCTTCACGGCGATTGCCTATCCGATGATCGGACTGCGGGCCGGAGTGCTGCACTTCTTCAACTGCCTGGCGCTGGTCACTCTGGTGGCCAATGTGTCAACGTCCTTCGGATATCTAATATCCTGCGCCAGCTCCTCGACCTCGATGGCGCTGTCTGTGGGTCCGCCGGTTATCATACCATTCCTGCTCTTTGGCGGCTTCTTCTTGAACTCGGGCTCGGTGCCAGTATACCTCAAATGGTTGTCGTACCTCTCATGGTTCCGTTACGCCAACGAGGGTCTGCTGATTAACCAATGGGCGGACGTGGAGCCGGGCGAAATTAGCTGCACATCGTCGAACACCACGTGCCCCAGTTCGGGCAAGGTCATCCTGGAGACGCTTAACTTCTCCGCCGCCGATCTGCCGCTGGACTACGTGGGTCTGGCCATTCTCATCGTGAGCTTCCGGGTGCTCGCATATCTGGCTCTAAGACTTCGGGCCCGACGCAAGGAGTAGCCGACATATATCCGAAATAACTGCTTGTTTTTTTTTTTTACCATTATTACCATCGTGTTTACTGTTTATTGCCCCCTCAAAAAGCTAATGTAATTATATTTGTGCCAATAAAAACAAGATATGACCTATAGAATACAAGTATTTCCCCTTCGAACATCCCCACAAGTAGACTTTGGATTTGTCTTCTAACCAAAAGACTTACACACCTGCATACCTTACATCAAAAACTCGTTTATCGCTACATAAAACACCGGGATATATTTTTTATATACATACTTTTCAAATCGCGCGCCCTCTTCATAATTCACCTCCACCACACCACGTTTCGTAGTTGCTCTTTCGCTGTCTCCCACCCGCTCTCCGCAACACATTCACCTTTTGTTCGACGACCTTGGAGCGACTGTCGTTAGTTCCGCGCGATTCGGTTCGCTCAAATGGTTCCGAGTGGTTCATTTCGTCTCAATAGAAATTAGTAATAAATATTTGTATGTACAATTTATTTGCTCCAATATATTTGTATATATTTCCCTCACAGCTATATTTATTCTAATTTAATATTATGACTTTTTAAGGTAATTTTTTGTGACCTGTTCGGAGTGATTAGCGTTACAATTTGAACTGAAAGTGACATCCAGTGTTTGTTCCTTGTGTAGATGCATCTCAAAAAAATGGTGGGCATAATAGTGTTGTTTATATATATCAAAAATAACAACTATAATAATAAGAATACATTTAATTTAGAAAATGCTTGGATTTCACTGGAACTAGGCTAGCATAACTTCGTATAATGTATGCTATACGAAGTTATGCTAGCGGATCCGGGAATTGGGAATTCACGTAAGTACTGTCTGCAGCGTAAGCTTCGTACGTAGCGACCGTCTCAAAGTACTGCCTTTCTGCGTTGGAAAACATCGCCTTTTTCGTCCAAAAGGAGTCCCCAGGTTCGATCCGCATGGCGTTGTGCGTGCGTGCCTTTCTTTTCAAATGATTACGGCTATTAACTTGGGGGCGTTAAGTTGGAAACACGTAAATTGCAGACTGCGATTAGAGTGACCATGAGTAGGAGTTCAAAATCTCCTGACATCATTTTCTTAAAACCTGCTTTGTTTTTTACATTTCTATTTAATATAACTCCTATTTGAATAAAAAAACAAAACAAGTTTAGATGTTAAGATATTAACTACATCCTTTGCTCCAAAGGGAGAGGGGAAGTTATGGAGTTAATTAATTTGCTGTTGGAAATCAATATGGAGTCAGAAATATAATGATTTACTAAACCTTATTGAATCGGTAACGATGCGAATTTATATTAAAATAGCTTTTATGAAACATTCAACAAAAATATTATTAATGTTGGCCCACTTTAGCAACCGGTTAGGTCTACCGGTTGGGCAAGCAAAGATTCACGCCCTGGTTCGAGTCCCAACTAGTCCTGCAAAATACCGCAGCAAGTTTTAGAGAGACCAAGTGCCATTACCTCTCCCACTTCAGTTATCGGTTATGCGGCGTTTAAGTCGACAGCTTGCCGTCTCTAGCTCCGGTGCCTATATAAAGCAGCCCGCTTTCCACATTTCATATTCGTTTTACGTTTGTCAAGCCTCATAGCCGGCAGTTCGAACGTATACGCTCTCTGAGTCAGACCTCGAAATCGTAGCTCTACACAATTCTGTGAATTTTCCTTGTCGCGTGTGAAACACTTCCAAGGCAATCTTACAAAATGAGTTTGCAAGACGAAGCTCTGAAGGCTTGGCAGGCTCGATACGGAGAACCAGCCAATCTCCCTGCTGCAGACACAGTCATCGCCCAGATGCTGCAGCATCGTTCCGTGCGCGCGTATTCGGACCTGCCGGTGGATGAACAAATGCTGAGTTGGGCGATTGCAGCAGCGCAAAGTGCCTCCACGAGTAGCAACCTGCAAGCATGGAGTGTCTTGGCAGTCCGCGACCGGGAGCGGTTGGCACGACTCGCCCGACTCTCGGGTAATCAGAGGCACGTCGAGCAGGCTCCTCTGTTCCTCGTCTGGTTGGTTGACTGGTCCCGTCTGAGGCGTTTGGCCCGTACCCTCCAAGCACCGACCGCGGGCATCGACTATCTGGAGTCGTATACAGTGGGAGTAGTAGATGCCGCACTGGCTGCTCAGAACGCGGCACTGGCTTTCGAAGCCCAAGGACTGGGTATCGTGTACATAGGTGGAATGAGGAATCACCCAGAAGCAATGAGCGAGGAGCTGGGATTGCCCAATGATACGTTCGCGGTTTTTGGAATGTGCGTGGGTCATCCTGACCCTGCTCAACCAGCGGAAATTAAGCCCAGGTTGGCGCAGAGTGTGGTCCTGCACCGGGAGAGGTATGAAGCTACTGAAGCTGAAGCAGTTAGTGTGGCAGCCTACGACCGTAGGATGTCGGATTTCCAGCACCGTCAACAGCGAGAGAATAGGTCGTGGTCCAGCCAGGCAGTTGAACGGGTGAAAGGCGCGGATTCCCTCAGCGGTCGTCACCGTTTGCGCGATGCGTTGAACACCCTGGGATTCGGACTCCGATAAGCCCGACTCTAGATCATAATCAGCCATACCACATTTGTAGAGGTTTTACTTGCTTTAAAAAACCTCCCACACCTCCCCCTGAACCTGAAACATAAAATGAATGCAATTGTTGTTGTTAACTTGTTTATTGCAGCTTATAATGGTTACAAATAAAGCAATAGCATCACAAATTTCACAAATAAAGCATTTTTTTCACTGCATTCTAGTTGTGGTTTGCCCAAACTCATCAATGTATCTTAGCGGCTCGAGGGTACCTCTAGAGATCCACTAGTGTCGACGATGTAGGTCACGGTCTCGAAGCCGCGGTGCGGGTGCCAGGGCGTGCCCTTGGGCTCCCCGGGCGCGTACTCCACCTCACCCATCTGGTCCATCATGATGAACGGGTCGAGGTGGCGGTAGTTGATCCCGGCGAACGCGCGGCGCACCGGGAAGCCCTCGCCCTCGAAACCGCTGGGCGCGGTGGTCACGGTGAGCACGGGACGTGCGACGGCGTCGGCGGGTGCGGATACGCGGGGCAGCGTCAGCGGGTTCTCGACGGTCACGGCGGGCATGTCGACACTAGTTCTAGCCAGCTTTTGTTCCCTTTAGTGAGGGTTAATTTCGAGCTTGGCGTAATCATGGTCATAGCTGTTTCCTGTGTGAAATTGTTATCCGCTCACAATTCCACACAACATACGAGCCGGAAGCATAAAGTGTAAAGCCTGGGGTGCCTAATGAGTGAGCTAACTCACATTAATTGCGTTGCGCTCACTGCCCGCTTTCCAGTCGGGAAACCTGTCGTGCCAGCTGCATTAATGAATCGGCCAACGCGCGGGGAGAGGCGGTTTGCGTATTGGGCGCTCTTCCGCTTCCTCGCTCACTGACTCGCTGCGCTCGGTCGTTCGGCTGCGGCGAGCGGTATCAGCTCACTCAAAGGCGGTAATACGGTTATCCACAGAATCAGGGGATAACGCAGGAAAGAACATGTGAGCAAAAGGCCAGCAAAAGGCCAGGAACCGTAAAAAGGCCGCGTTGCTGGCGTTTTTCCATAGGCTCCGCCCCCCTGACGAGCATCACAAAAATCGACGCTCAAGTCAGAGGTGGCGAAACCCGACAGGACTATAAAGATACCAGGCGTTTCCCCCTGGAAGCTCCCTCGTGCGCTCTCCTGTTCCGACCCTGCCGCTTACCGGATACCTGTCCGCCTTTCTCCCTTCGGGAAGCGTGGCGCTTTCTCATAGCTCACGCTGTAGGTATCTCAGTTCGGTGTAGGTCGTTCGCTCCAAGCTGGGCTGTGTGCACGAACCCCCCGTTCAGCCCGACCGCTGCGCCTTATCCGGTAACTATCGTCTTGAGTCCAACCCGGTAAGACACGACTTATCGCCACTGGCAGCAGCCACTGGTAACAGGATTAGCAGAGCGAGGTATGTAGGCGGTGCTACAGAGTTCTTGAAGTGGTGGCCTAACTACGGCTACACTAGAAGAACAGTATTTGGTATCTGCGCTCTGCTGAAGCCAGTTACCTTCGGAAAAAGAGTTGGTAGCTCTTGATCCGGCAAACAAACCACCGCTGGTAGCGGTGGTTTTTTTGTTTGCAAGCAGCAGATTACGCGCAGAAAAAAAGGATCTCAAGAAGATCCTTTGATCTTTTCTACGGGGTCTGACGCTCAGTGGAACGAAAACTCACGTTAAGGGATTTTGGTCATGAGATTATCAAAAAGGATCTTCACCTAGATCCTTTTAAATTAAAAATGAAGTTTTAAATCAATCTAAAGTATATATGAGTAAACTTGGTCTGACAGTTACCAATGCTTAATCAGTGAGGCACCTATCTCAGCGATCTGTCTATTTCGTTCATCCATAGTTGCCTGACTCCCCGTCGTGTAGATAACTACGATACGGGAGGGCTTACCATCTGGCCCCAGTGCTGCAATGATACCGCGAGACCCACGCTCACCGGCTCCAGATTTATCAGCAATAAACCAGCCAGCCGGAAGGGCCGAGCGCAGAAGTGGTCCTGCAACTTTATCCGCCTCCATCCAGTCTATTAATTGTTGCCGGGAAGCTAGAGTAAGTAGTTCGCCAGTTAATAGTTTGCGCAACGTTGTTGCCATTGCTACAGGCATCGTGGTGTCACGCTCGTCGTTTGGTATGGCTTCATTCAGCTCCGGTTCCCAACGATCAAGGCGAGTTACATGATCCCCCATGTTGTGCAAAAAAGCGGTTAGCTCCTTCGGTCCTCCGATCGTTGTCAGAAGTAAGTTGGCCGCAGTGTTATCACTCATGGTTATGGCAGCACTGCATAATTCTCTTACTGTCATGCCATCCGTAAGATGCTTTTCTGTGACTGGTGAGTACTCAACCAAGTCATTCTGAGAATAGTGTATGCGGCGACCGAGTTGCTCTTGCCCGGCGTCAATACGGGATAATACCGCGCCACATAGCAGAACTTTAAAAGTGCTCATCATTGGAAAACGTTCTTCGGGGCGAAAACTCTCAAGGATCTTACCGCTGTTGAGATCCAGTTCGATGTAACCCACTCGTGCACCCAACTGATCTTCAGCATCTTTTACTTTCACCAGCGTTTCTGGGTGAGCAAAAACAGGAAGGCAAAATGCCGCAAAAAAGGGAATAAGGGCGACACGGAAATGTTGAATACTCATACTCTTCCTTTTTCAATATTATTGAAGCATTTATCAGGGTTATTGTCTCATGAGCGGATACATATTTGAATGTATTTAGAAAAATAAACAAATAGGGGTTCCGCGCACATTTCCTCCGAAAAGTGCCACCTAAATTGTAAGCGTTAATATTTTTGTTAAAATTCGCGTTAAATTTTTTGTTAAATCAGCTCATTTTTTTAACCAATAGGCCGAAATCGGCAAAAATCCCCTTATAAATCAAAAAGAATAGACCGAGATAGGGTTGAGTGTTGTTCCCAGTTTTGGC |
| pKT111.SnrA | AAAATAATCAATTAGGGGTTCCGCGCAACAATTTCCCCGAAAAAAGTGCCCACCTAAATTGTAAAGCGTTTAATATTTTTGTTTAAAATTCGCGTTAAATTTTTTGTTAAATCAGCTCATTTTTTTAACCAATAGGCCGAAAATCGGCAAAAATCCCTTATAAATCAAAAAGAATAGACCGAGATAGGGTTGAGTGTTGTTCCAGTTTGGAACAAGAGTCCACTATTAAAGAACGTGGACTCCAACGTCAAAGGGCGAAAAACCGTCTATCAGGGCGATGGCCCACTACGTGAACCATCACCCTAATCAAGTTTTTTGGGGTCGAGGTGCCGTAAAGCACTAAATCGGAACCCTAAAGGGAGCCCCCGATTTAGAGCTTGACGGGGAAAGCCGGCGAACGTGGCGAGAAAGGAAGGGAAGAAAGCGAAAGGAGCGGGCGCTAGGGCGCTGGCAAGTGTAGCGGTCACGCTGCGCGTAACCACCACACCCGCCGCGCTTAATGCGCCGCTACAGGGCGCGTCCCATTCGCCATTCAGGCTGCGCAACTGTTGGGAAGGGCGATCGGTGCGGGCCTCTTCGCTATTACGCCAGCTGGCGAAAGGGGGATGTGCTGCAAGGCGATTAAGTTGGGTAACGCCAGGGTTTTCCCAGTCACGACGTTGTAAAACGACGGCCAGTGAATTGTAATACGACTCACTATAGGGCGAATTGGGTACGTACCGGGCCCCTAGTATGTATGTAAGTTAATAAAACCCATTTTTGCGGAAAGTAGATAAAAAAAACATTTTTTTTTTTTACTGCACTGGATATCATTGAACTTATCTGATCAGTTTTAAATTTACTTCGATCCAAGGGTATTTGATGTACCAGGTTCTTTCGATTACCTCTCACTCAAAATGACATTCCACTCAAAGTCAGCGCTGTTTGCCTCCTTCTCTGTCCACAGAAATATCGCCGTCTCTTTCGCCGCTGCGTCCGCTATCTCTTTCGCCACCGTTTGTAGCGTTACGTAGCGTCAATGTCCGCCTTCAGTTGCATTTTGTCAGCGGTTTCGTGACGAAGCTCCAAGCGGTTTACGCCATCAATTAAACACAAAGTGCTGTGCCAAAACTCCTCTCGCTTCTTATTTTTGTTTGTTTTTTGAGTGATTGGGGTGGTGATTGGTTTTGGGTGGGTAAGCAGGGGAAAGTGTGAAAAATCCCGGCAATGGGCCAAGAGGATCAGGAGCTATTAATTCGCGGAGGCAGCAAACACCCATCTGCCGAGCATCTGAACAATGTGAGTAGTACATGTGCATACATCTTAAGTTCACTTGATCTATAGGAACTGCGATTGCAACATCAAATTGTCTGCGGCGTGAGAACTGCGACCCACAAAAATCCCAAACCGCAATTGCACAAACAAATAGTGACACGAAACAGATTATTCTGGTAGCTGTTCTCGCTATATAAGACAATTTTTGAGATCATATCATGATCAAGACATCTAAAGGCATTCATTTTCGACTATATTCTTTTTTACAAAAAATATAACAACCAGATATTTTAAGCTGATCCTAGATGCACAAAAAATAAATAAAAGTATAAACCTACTTCGTAGGATACTTCGGGGTACTTTTTGTTCGGGGTTAGATGAGCATAACGCTTGTAGTTGATATTTGAGATCCCCTATCATTGCAGGGTGACAGCGGAGCGGCTTCGCAGAGCTGCATTAACCAGGGCTTCGGGCAGGCCAAAAACTACGGCACGCTCCGGCCACCCAGTCCGCCGGAGGACTCCGGTTCAGGGAGCGGCCAACTAGCCGAGAACCTCACCTATGCCTGGCACAATATGGACATCTTTGGGGCGGTCAATCAGCCGGGCTCCGGATGGCGGCAGCTGGTCAACCGGACACGCGGACTATTCTGCAACGAGCGACACATACCGGCGCCCAGGAAACATTTGCTCAAGAACGGTGAGTTTCTATTCGCAGTCGGCTGATCTGTGTGAAATCTTAATAAAGGGTCCAATTACCAATTTGAAACTCAGTTTGCGGCGTGGCCTATCCGGGCGAACTTTTGGCCGTGATGGGCAGTTCCGGTGCCGGAAAGACGACCCTGCTGAATGCCCTTGCCTTTCGATCGCCGCAGGGCATCCAAGTATCGCCATCCGGGATGCGACTGCTCAATGGCCAACCTGTGGACGCCAAGGAGATGCAGGCCAGGTGCGCCTATGTCCAGCAGGATGACCTCTTTATCGGCTCCCTAACGGCCAGGGAACACCTGATTTTCCAAGCCATGGTGCGGATGCCACGACATCTGACCTATCGGCAGCGAGTGGCCCGCGTGGATCAGGTGATCCAGGAGCTTTCGCTCAGCAAATGTCAGCACACGATCATCGGTGTGCCCGGCAGGGTGAAAGGTCTGTCCGGCGGAGAAAGGAAGCGTCTGGCATTCGCCTCCGAGGCTCTAACCGATCCGCCGCTTCTGATCTGCGATGAGCCCACCTCCGGACTGGACTCCTTTACCGCCCACAGCGTCGTCCAGGTGCTGAAGAAGCTGTCGCAGAAGGGCAAGACCGTCATCCTGACCATTCATCAGCCGTCTTCCGAGCTGTTTGAGCTCTTTGACAAGATCCTTCTGATGGCCGAGGGCAGGGTAGCTTTCTTGGGCACTCCCAGCGAAGCCGTCGACTTCTTTTCCTAGTGAGTTCGATGTGTTTATTAAGGGTATCTAGTATTACATAACATCTCAACTCCTATCCAGCGTGGGTGCCCAGTGTCCTACCAACTACAATCCGGCGGACTTTTACGTACAGGTGTTGGCCGTTGTGCCCGGACGGGAGATCGAGTCCCGTGATCGGATCGCCAAGATATGCGACAATTTTGCCATTAGCAAAGTAGCCCGGGATATGGAGCAGTTGTTGGCCACCAAAAATCTGGAGAAGCCACTGGAGCAGCCGGAGAATGGGTACACCTACAAGGCCACCTGGTTCATGCAGTTCCGGGCGGTCCTGTGGCGATCCTGGCTGTCGGTGCTCAAGGAACCACTCCTCGTAAAAGTGCGACTTATTCAGACAACGGTGAGTGGTTCCAGTGGAAACAAATGATATAACGCTTACAATTCTTGGAAACAAATTCGCTAGATTTTAGATAGAATTGCCTGATTCCACACCCTTCTTAGTTTTTTTCAATGAGATGTATAGTTTATAGTTTTGCAGAAGATAAATAAATTTCATTTAACTCGCGAATATTAATGAGATGCGAGTAACATTTTAATTTGCAGATGGTTGCCATCTTGATTGGCCTCATCTTTTTGGGCCAACAACTCACGCAAGTGGGTGTGATGAATATCAACGGAGCCATCTTCCTCTTCCTGACCAACATGACCTTTCAAAACGTCTTTGCCACGATAAATGTAAGTCATGTTTAGAATACATTTGCATTTCAATAATTTACTAACTTTCTAATGAATCGATTCGATTTAGGTGTTCACCTCAGAGCTGCCAGTTTTTATGAGGGAGGCCCGAAGTCGACTTTATCGCTGTGACACATACTTTCTGGGCAAAACGATTGCCGAATTGCCGCTTTTTCTCACAGTGCCACTGGTCTTCACGGCGATTGCCTATCCGATGATCGGACTGCGGGCCGGAGTGCTGCACTTCTTCAACTGCCTGGCGCTGGTCACTCTGGTGGCCAATGTGTCAACGTCCTTCGGATATCTAATATCCTGCGCCAGCTCCTCGACCTCGATGGCGCTGTCTGTGGGTCCGCCGGTTATCATACCATTCCTGCTCTTTGGCGGCTTCTTCTTGAACTCGGGCTCGGTGCCAGTATACCTCAAATGGTTGTCGTACCTCTCATGGTTCCGTTACGCCAACGAGGGTCTGCTGATTAACCAATGGGCGGACGTGGAGCCGGGCGAAATTAGCTGCACATCGTCGAACACCACGTGCCCCAGTTCGGGCAAGGTCATCCTGGAGACGCTTAACTTCTCCGCCGCCGATCTGCCGCTGGACTACGTGGGTCTGGCCATTCTCATCGTGAGCTTCCGGGTGCTCGCATATCTGGCTCTAAGACTTCGGGCCCGACGCAAGGAGTAGCCGACATATATCCGAAATAACTGCTTGTTTTTTTTTTTTACCATTATTACCATCGTGTTTACTGTTTATTGCCCCCTCAAAAAGCTAATGTAATTATATTTGTGCCAATAAAAACAAGATATGACCTATAGAATACAAGTATTTCCCCTTCGAACATCCCCACAAGTAGACTTTGGATTTGTCTTCTAACCAAAAGACTTACACACCTGCATACCTTACATCAAAAACTCGTTTATCGCTACATAAAACACCGGGATATATTTTTTATATACATACTTTTCAAATCGCGCGCCCTCTTCATAATTCACCTCCACCACACCACGTTTCGTAGTTGCTCTTTCGCTGTCTCCCACCCGCTCTCCGCAACACATTCACCTTTTGTTCGACGACCTTGGAGCGACTGTCGTTAGTTCCGCGCGATTCGGTTCGCTCAAATGGTTCCGAGTGGTTCATTTCGTCTCAATAGAAATTAGTAATAAATATTTGTATGTACAATTTATTTGCTCCAATATATTTGTATATATTTCCCTCACAGCTATATTTATTCTAATTTAATATTATGACTTTTTAAGGTAATTTTTTGTGACCTGTTCGGAGTGATTAGCGTTACAATTTGAACTGAAAGTGACATCCAGTGTTTGTTCCTTGTGTAGATGCATCTCAAAAAAATGGTGGGCATAATAGTGTTGTTTATATATATCAAAAATAACAACTATAATAATAAGAATACATTTAATTTAGAAAATGCTTGGATTTCACTGGAACTAGGCTAGCATAACTTCGTATAATGTATGCTATACGAAGTTATGCTAGCGGATCCGGGAATTGGGAATTCACGTAAGTACTGTCTGCAGCGTAAGCTTCGTACGTAGCGACCGTCTCAAAGTACTGCCTTTCTGCGTTGGAAAACATCGCCTTTTTCGTCCAAAAGGAGTCCCCAGGTTCGATCCGCATGGCGTTGTGCGTGCGTGCCTTTCTTTTCAAATGATTACGGCTATTAACTTGGGGGCGTTAAGTTGGAAACACGTAAATTGCAGACTGCGATTAGAGTGACCATGAGTAGGAGTTCAAAATCTCCTGACATCATTTTCTTAAAACCTGCTTTGTTTTTTACATTTCTATTTAATATAACTCCTATTTGAATAAAAAAACAAAACAAGTTTAGATGTTAAGATATTAACTACATCCTTTGCTCCAAAGGGAGAGGGGAAGTTATGGAGTTAATTAATTTGCTGTTGGAAATCAATATGGAGTCAGAAATATAATGATTTACTAAACCTTATTGAATCGGTAACGATGCGAATTTATATTAAAATAGCTTTTATGAAACATTCAACAAAAATATTATTAATGTTGGCCCACTTTAGCAACCGGTTAGGTCTACCGGTTGGGCAAGCAAAGATTCACGCCCTGGTTCGAGTCCCAACTAGTCCTGCAAAATACCGCAGCAAGTTTTAGAGAGACCAAGTGCCATTACCTCTCCCACTTCAGTTATCGGTTATGCGGCGTTTAAGTCGACAGCTTGCCGTCTCTAGCTCCGGTGCCTATATAAAGCAGCCCGCTTTCCACATTTCATATTCGTTTTACGTTTGTCAAGCCTCATAGCCGGCAGTTCGAACGTATACGCTCTCTGAGTCAGACCTCGAAATCGTAGCTCTACACAATTCTGTGAATTTTCCTTGTCGCGTGTGAAACACTTCCAAGGCAATCTTACAAAATGAGTCCAACTATAGAACTCCTGTGCGGACATCGAAGTATTCGTCATTTTACCGACGAGCCTGTAACGGAGCCACAGCGCGAAGCCATCAACTGCCGGGCTCGCTCGACTTCGAGTTCGAGTTTCTTGCAATGTAGCAGCATAATCCGTATCACTGATCGTGCGTTGCGAGAGGCTCTCTCCTGCCTGACGGGCGGACAAAAGCATGTTGCACAGGCCGCCGAGTTCTGGGTTTTTTGCGCCGATTTTAATAGGCATCTGCAGATATGCCCCGACGCTCAGCTGGGACTGGCCGAACAGTTGCTCTTGGGAGTCGTGGACACTGCCATGATGGGTCAAAATGCGCTGACGGCTGCTGAATCCTTGGGACTGGGAGGCGTATATATCGGTGGCATCCGCAACAACATCGAAAGCGTCACTGAACTGCTGAAGCTGCCCAAACACGTTCTGCCCCTGTTTGGTTTGTGTCTGGGCTGGCCCGCCGGTAATCCGGATCTGAAACCGAGGTTGCCCGCCGAACTGGTTGTGCACGAGAATCAGTATCAACCGCTCGATGAGAAGCTGCTCGCTCGCTATGATGAGCAACTGGCTGAATACTACCTGACGCGTGGCAGTAACACCCGGCGTGATACGTGGTCCGACCATATTCGTAGGACTCTCATAAAAGAGAATCGACCATTCATTTTGGAATATTTGCATAAACAAGGCTGGGCTACCCGATAAGCCCGACTCTAGATCATAATCAGCCATACCACATTTGTAGAGGTTTTACTTGCTTTAAAAAACCTCCCACACCTCCCCCTGAACCTGAAACATAAAATGAATGCAATTGTTGTTGTTAACTTGTTTATTGCAGCTTATAATGGTTACAAATAAAGCAATAGCATCACAAATTTCACAAATAAAGCATTTTTTTCACTGCATTCTAGTTGTGGTTTGCCCAAACTCATCAATGTATCTTAGCGGCTCGAGGGTACCTCTAGAGATCCACTAGTGTCGACGATGTAGGTCACGGTCTCGAAGCCGCGGTGCGGGTGCCAGGGCGTGCCCTTGGGCTCCCCGGGCGCGTACTCCACCTCACCCATCTGGTCCATCATGATGAACGGGTCGAGGTGGCGGTAGTTGATCCCGGCGAACGCGCGGCGCACCGGGAAGCCCTCGCCCTCGAAACCGCTGGGCGCGGTGGTCACGGTGAGCACGGGACGTGCGACGGCGTCGGCGGGTGCGGATACGCGGGGCAGCGTCAGCGGGTTCTCGACGGTCACGGCGGGCATGTCGACACTAGTTCTAGCCAGCTTTTGTTCCCTTTAGTGAGGGTTAATTTCGAGCTTGGCGTAATCATGGTCATAGCTGTTTCCTGTGTGAAATTGTTATCCGCTCACAATTCCACACAACATACGAGCCGGAAGCATAAAGTGTAAAGCCTGGGGTGCCTAATGAGTGAGCTAACTCACATTAATTGCGTTGCGCTCACTGCCCGCTTTCCAGTCGGGAAACCTGTCGTGCCAGCTGCATTAATGAATCGGCCAACGCGCGGGGAGAGGCGGTTTGCGTATTGGGCGCTCTTCCGCTTCCTCGCTCACTGACTCGCTGCGCTCGGTCGTTCGGCTGCGGCGAGCGGTATCAGCTCACTCAAAGGCGGTAATACGGTTATCCACAGAATCAGGGGATAACGCAGGAAAGAACATGTGAGCAAAAGGCCAGCAAAAGGCCAGGAACCGTAAAAAGGCCGCGTTGCTGGCGTTTTTCCATAGGCTCCGCCCCCCTGACGAGCATCACAAAAATCGACGCTCAAGTCAGAGGTGGCGAAACCCGACAGGACTATAAAGATACCAGGCGTTTCCCCCTGGAAGCTCCCTCGTGCGCTCTCCTGTTCCGACCCTGCCGCTTACCGGATACCTGTCCGCCTTTCTCCCTTCGGGAAGCGTGGCGCTTTCTCATAGCTCACGCTGTAGGTATCTCAGTTCGGTGTAGGTCGTTCGCTCCAAGCTGGGCTGTGTGCACGAACCCCCCGTTCAGCCCGACCGCTGCGCCTTATCCGGTAACTATCGTCTTGAGTCCAACCCGGTAAGACACGACTTATCGCCACTGGCAGCAGCCACTGGTAACAGGATTAGCAGAGCGAGGTATGTAGGCGGTGCTACAGAGTTCTTGAAGTGGTGGCCTAACTACGGCTACACTAGAAGAACAGTATTTGGTATCTGCGCTCTGCTGAAGCCAGTTACCTTCGGAAAAAGAGTTGGTAGCTCTTGATCCGGCAAACAAACCACCGCTGGTAGCGGTGGTTTTTTTGTTTGCAAGCAGCAGATTACGCGCAGAAAAAAAGGATCTCAAGAAGATCCTTTGATCTTTTCTACGGGGTCTGACGCTCAGTGGAACGAAAACTCACGTTAAGGGATTTTGGTCATGAGATTATCAAAAAGGATCTTCACCTAGATCCTTTTAAATTAAAAATGAAGTTTTAAATCAATCTAAAGTATATATGAGTAAACTTGGTCTGACAGTTACCAATGCTTAATCAGTGAGGCACCTATCTCAGCGATCTGTCTATTTCGTTCATCCATAGTTGCCTGACTCCCCGTCGTGTAGATAACTACGATACGGGAGGGCTTACCATCTGGCCCCAGTGCTGCAATGATACCGCGAGACCCACGCTCACCGGCTCCAGATTTATCAGCAATAAACCAGCCAGCCGGAAGGGCCGAGCGCAGAAGTGGTCCTGCAACTTTATCCGCCTCCATCCAGTCTATTAATTGTTGCCGGGAAGCTAGAGTAAGTAGTTCGCCAGTTAATAGTTTGCGCAACGTTGTTGCCATTGCTACAGGCATCGTGGTGTCACGCTCGTCGTTTGGTATGGCTTCATTCAGCTCCGGTTCCCAACGATCAAGGCGAGTTACATGATCCCCCATGTTGTGCAAAAAAGCGGTTAGCTCCTTCGGTCCTCCGATCGTTGTCAGAAGTAAGTTGGCCGCAGTGTTATCACTCATGGTTATGGCAGCACTGCATAATTCTCTTACTGTCATGCCATCCGTAAGATGCTTTTCTGTGACTGGTGAGTACTCAACCAAGTCATTCTGAGAATAGTGTATGCGGCGACCGAGTTGCTCTTGCCCGGCGTCAATACGGGATAATACCGCGCCACATAGCAGAACTTTAAAAGTGCTCATCATTGGAAAACGTTCTTCGGGGCGAAAACTCTCAAGGATCTTACCGCTGTTGAGATCCAGTTCGATGTAACCCACTCGTGCACCCAACTGATCTTCAGCATCTTTTACTTTCACCAGCGTTTCTGGGTGAGCAAAAACAGGAAGGCAAAATGCCGCAAAAAAGGGAATAAGGGCGACACGGAAATGTTGAATACTCATACTCTTCCTTTTTCAATATTATTGAAGCATTTATCAGGGTTATTGTCTCATGAGCGGATACATATTTGAATGTATTTAGAAAAATAAACAAATAGGGGTTCCGCGCACATTTCCTCCGAAAAGTGCCACCTAAATTGTAAGCGTTAATATTTTTGTTAAAATTCGCGTTAAATTTTTTGTTAAATCAGCTCATTTTTTTAACCAATAGGCCGAAATCGGCAAAAATCCCCTTATAAATCAAAAAGAATAGACCGAGATAGGGTTGAGTGTTGTTCCCAGTTTTGGC |
| pKT111.Mm-nr | AAAATAATCAATTAGGGGTTCCGCGCAACAATTTCCCCGAAAAAAGTGCCCACCTAAATTGTAAAGCGTTTAATATTTTTGTTTAAAATTCGCGTTAAATTTTTTGTTAAATCAGCTCATTTTTTTAACCAATAGGCCGAAAATCGGCAAAAATCCCTTATAAATCAAAAAGAATAGACCGAGATAGGGTTGAGTGTTGTTCCAGTTTGGAACAAGAGTCCACTATTAAAGAACGTGGACTCCAACGTCAAAGGGCGAAAAACCGTCTATCAGGGCGATGGCCCACTACGTGAACCATCACCCTAATCAAGTTTTTTGGGGTCGAGGTGCCGTAAAGCACTAAATCGGAACCCTAAAGGGAGCCCCCGATTTAGAGCTTGACGGGGAAAGCCGGCGAACGTGGCGAGAAAGGAAGGGAAGAAAGCGAAAGGAGCGGGCGCTAGGGCGCTGGCAAGTGTAGCGGTCACGCTGCGCGTAACCACCACACCCGCCGCGCTTAATGCGCCGCTACAGGGCGCGTCCCATTCGCCATTCAGGCTGCGCAACTGTTGGGAAGGGCGATCGGTGCGGGCCTCTTCGCTATTACGCCAGCTGGCGAAAGGGGGATGTGCTGCAAGGCGATTAAGTTGGGTAACGCCAGGGTTTTCCCAGTCACGACGTTGTAAAACGACGGCCAGTGAATTGTAATACGACTCACTATAGGGCGAATTGGGTACGTACCGGGCCCCTAGTATGTATGTAAGTTAATAAAACCCATTTTTGCGGAAAGTAGATAAAAAAAACATTTTTTTTTTTTACTGCACTGGATATCATTGAACTTATCTGATCAGTTTTAAATTTACTTCGATCCAAGGGTATTTGATGTACCAGGTTCTTTCGATTACCTCTCACTCAAAATGACATTCCACTCAAAGTCAGCGCTGTTTGCCTCCTTCTCTGTCCACAGAAATATCGCCGTCTCTTTCGCCGCTGCGTCCGCTATCTCTTTCGCCACCGTTTGTAGCGTTACGTAGCGTCAATGTCCGCCTTCAGTTGCATTTTGTCAGCGGTTTCGTGACGAAGCTCCAAGCGGTTTACGCCATCAATTAAACACAAAGTGCTGTGCCAAAACTCCTCTCGCTTCTTATTTTTGTTTGTTTTTTGAGTGATTGGGGTGGTGATTGGTTTTGGGTGGGTAAGCAGGGGAAAGTGTGAAAAATCCCGGCAATGGGCCAAGAGGATCAGGAGCTATTAATTCGCGGAGGCAGCAAACACCCATCTGCCGAGCATCTGAACAATGTGAGTAGTACATGTGCATACATCTTAAGTTCACTTGATCTATAGGAACTGCGATTGCAACATCAAATTGTCTGCGGCGTGAGAACTGCGACCCACAAAAATCCCAAACCGCAATTGCACAAACAAATAGTGACACGAAACAGATTATTCTGGTAGCTGTTCTCGCTATATAAGACAATTTTTGAGATCATATCATGATCAAGACATCTAAAGGCATTCATTTTCGACTATATTCTTTTTTACAAAAAATATAACAACCAGATATTTTAAGCTGATCCTAGATGCACAAAAAATAAATAAAAGTATAAACCTACTTCGTAGGATACTTCGGGGTACTTTTTGTTCGGGGTTAGATGAGCATAACGCTTGTAGTTGATATTTGAGATCCCCTATCATTGCAGGGTGACAGCGGAGCGGCTTCGCAGAGCTGCATTAACCAGGGCTTCGGGCAGGCCAAAAACTACGGCACGCTCCGGCCACCCAGTCCGCCGGAGGACTCCGGTTCAGGGAGCGGCCAACTAGCCGAGAACCTCACCTATGCCTGGCACAATATGGACATCTTTGGGGCGGTCAATCAGCCGGGCTCCGGATGGCGGCAGCTGGTCAACCGGACACGCGGACTATTCTGCAACGAGCGACACATACCGGCGCCCAGGAAACATTTGCTCAAGAACGGTGAGTTTCTATTCGCAGTCGGCTGATCTGTGTGAAATCTTAATAAAGGGTCCAATTACCAATTTGAAACTCAGTTTGCGGCGTGGCCTATCCGGGCGAACTTTTGGCCGTGATGGGCAGTTCCGGTGCCGGAAAGACGACCCTGCTGAATGCCCTTGCCTTTCGATCGCCGCAGGGCATCCAAGTATCGCCATCCGGGATGCGACTGCTCAATGGCCAACCTGTGGACGCCAAGGAGATGCAGGCCAGGTGCGCCTATGTCCAGCAGGATGACCTCTTTATCGGCTCCCTAACGGCCAGGGAACACCTGATTTTCCAAGCCATGGTGCGGATGCCACGACATCTGACCTATCGGCAGCGAGTGGCCCGCGTGGATCAGGTGATCCAGGAGCTTTCGCTCAGCAAATGTCAGCACACGATCATCGGTGTGCCCGGCAGGGTGAAAGGTCTGTCCGGCGGAGAAAGGAAGCGTCTGGCATTCGCCTCCGAGGCTCTAACCGATCCGCCGCTTCTGATCTGCGATGAGCCCACCTCCGGACTGGACTCCTTTACCGCCCACAGCGTCGTCCAGGTGCTGAAGAAGCTGTCGCAGAAGGGCAAGACCGTCATCCTGACCATTCATCAGCCGTCTTCCGAGCTGTTTGAGCTCTTTGACAAGATCCTTCTGATGGCCGAGGGCAGGGTAGCTTTCTTGGGCACTCCCAGCGAAGCCGTCGACTTCTTTTCCTAGTGAGTTCGATGTGTTTATTAAGGGTATCTAGTATTACATAACATCTCAACTCCTATCCAGCGTGGGTGCCCAGTGTCCTACCAACTACAATCCGGCGGACTTTTACGTACAGGTGTTGGCCGTTGTGCCCGGACGGGAGATCGAGTCCCGTGATCGGATCGCCAAGATATGCGACAATTTTGCCATTAGCAAAGTAGCCCGGGATATGGAGCAGTTGTTGGCCACCAAAAATCTGGAGAAGCCACTGGAGCAGCCGGAGAATGGGTACACCTACAAGGCCACCTGGTTCATGCAGTTCCGGGCGGTCCTGTGGCGATCCTGGCTGTCGGTGCTCAAGGAACCACTCCTCGTAAAAGTGCGACTTATTCAGACAACGGTGAGTGGTTCCAGTGGAAACAAATGATATAACGCTTACAATTCTTGGAAACAAATTCGCTAGATTTTAGATAGAATTGCCTGATTCCACACCCTTCTTAGTTTTTTTCAATGAGATGTATAGTTTATAGTTTTGCAGAAGATAAATAAATTTCATTTAACTCGCGAATATTAATGAGATGCGAGTAACATTTTAATTTGCAGATGGTTGCCATCTTGATTGGCCTCATCTTTTTGGGCCAACAACTCACGCAAGTGGGTGTGATGAATATCAACGGAGCCATCTTCCTCTTCCTGACCAACATGACCTTTCAAAACGTCTTTGCCACGATAAATGTAAGTCATGTTTAGAATACATTTGCATTTCAATAATTTACTAACTTTCTAATGAATCGATTCGATTTAGGTGTTCACCTCAGAGCTGCCAGTTTTTATGAGGGAGGCCCGAAGTCGACTTTATCGCTGTGACACATACTTTCTGGGCAAAACGATTGCCGAATTGCCGCTTTTTCTCACAGTGCCACTGGTCTTCACGGCGATTGCCTATCCGATGATCGGACTGCGGGCCGGAGTGCTGCACTTCTTCAACTGCCTGGCGCTGGTCACTCTGGTGGCCAATGTGTCAACGTCCTTCGGATATCTAATATCCTGCGCCAGCTCCTCGACCTCGATGGCGCTGTCTGTGGGTCCGCCGGTTATCATACCATTCCTGCTCTTTGGCGGCTTCTTCTTGAACTCGGGCTCGGTGCCAGTATACCTCAAATGGTTGTCGTACCTCTCATGGTTCCGTTACGCCAACGAGGGTCTGCTGATTAACCAATGGGCGGACGTGGAGCCGGGCGAAATTAGCTGCACATCGTCGAACACCACGTGCCCCAGTTCGGGCAAGGTCATCCTGGAGACGCTTAACTTCTCCGCCGCCGATCTGCCGCTGGACTACGTGGGTCTGGCCATTCTCATCGTGAGCTTCCGGGTGCTCGCATATCTGGCTCTAAGACTTCGGGCCCGACGCAAGGAGTAGCCGACATATATCCGAAATAACTGCTTGTTTTTTTTTTTTACCATTATTACCATCGTGTTTACTGTTTATTGCCCCCTCAAAAAGCTAATGTAATTATATTTGTGCCAATAAAAACAAGATATGACCTATAGAATACAAGTATTTCCCCTTCGAACATCCCCACAAGTAGACTTTGGATTTGTCTTCTAACCAAAAGACTTACACACCTGCATACCTTACATCAAAAACTCGTTTATCGCTACATAAAACACCGGGATATATTTTTTATATACATACTTTTCAAATCGCGCGCCCTCTTCATAATTCACCTCCACCACACCACGTTTCGTAGTTGCTCTTTCGCTGTCTCCCACCCGCTCTCCGCAACACATTCACCTTTTGTTCGACGACCTTGGAGCGACTGTCGTTAGTTCCGCGCGATTCGGTTCGCTCAAATGGTTCCGAGTGGTTCATTTCGTCTCAATAGAAATTAGTAATAAATATTTGTATGTACAATTTATTTGCTCCAATATATTTGTATATATTTCCCTCACAGCTATATTTATTCTAATTTAATATTATGACTTTTTAAGGTAATTTTTTGTGACCTGTTCGGAGTGATTAGCGTTACAATTTGAACTGAAAGTGACATCCAGTGTTTGTTCCTTGTGTAGATGCATCTCAAAAAAATGGTGGGCATAATAGTGTTGTTTATATATATCAAAAATAACAACTATAATAATAAGAATACATTTAATTTAGAAAATGCTTGGATTTCACTGGAACTAGGCTAGCATAACTTCGTATAATGTATGCTATACGAAGTTATGCTAGCGGATCCGGGAATTGGGAATTCACGTAAGTACTGTCTGCAGCGTAAGCTTCGTACGTAGCGACCGTCTCAAAGTACTGCCTTTCTGCGTTGGAAAACATCGCCTTTTTCGTCCAAAAGGAGTCCCCAGGTTCGATCCGCATGGCGTTGTGCGTGCGTGCCTTTCTTTTCAAATGATTACGGCTATTAACTTGGGGGCGTTAAGTTGGAAACACGTAAATTGCAGACTGCGATTAGAGTGACCATGAGTAGGAGTTCAAAATCTCCTGACATCATTTTCTTAAAACCTGCTTTGTTTTTTACATTTCTATTTAATATAACTCCTATTTGAATAAAAAAACAAAACAAGTTTAGATGTTAAGATATTAACTACATCCTTTGCTCCAAAGGGAGAGGGGAAGTTATGGAGTTAATTAATTTGCTGTTGGAAATCAATATGGAGTCAGAAATATAATGATTTACTAAACCTTATTGAATCGGTAACGATGCGAATTTATATTAAAATAGCTTTTATGAAACATTCAACAAAAATATTATTAATGTTGGCCCACTTTAGCAACCGGTTAGGTCTACCGGTTGGGCAAGCAAAGATTCACGCCCTGGTTCGAGTCCCAACTAGTCCTGCAAAATACCGCAGCAAGTTTTAGAGAGACCAAGTGCCATTACCTCTCCCACTTCAGTTATCGGTTATGCGGCGTTTAAGTCGACAGCTTGCCGTCTCTAGCTCCGGTGCCTATATAAAGCAGCCCGCTTTCCACATTTCATATTCGTTTTACGTTTGTCAAGCCTCATAGCCGGCAGTTCGAACGTATACGCTCTCTGAGTCAGACCTCGAAATCGTAGCTCTACACAATTCTGTGAATTTTCCTTGTCGCGTGTGAAACACTTCCAAGGCAATCTTACAAAATGACCATTACCCAGGCTGTGATGCAGCGTTATTCCACAAAAAGTTTTGATCCGGCGAAGAAAATATCGGACGAGGTAATGAACTCGATTAAAACGCTCTTGCGGTACAGTCCAAGCTCCGTAAATATACAACCATGGCATTTTATAATCGCAGCCTCGGAAGAGGGTAAAAATAGGATCGCCGCCTCGACTCGGCCGGGATTCGAATTTAACACGGCCAAGATTACCGATGCAAGCCATGTAGTGTTGTTGTGTGCGAAAACGGATGTAGATGAGGCATATATGAATAGCGTACTGGAGCAAGAAGATAAAGATGGAAGGTACACTACCCCTGAACATAAGGCTATGAATAACGGAGGTCGAACCTTTTTTGTCAATTTGCACAAGGAAACTCTGGGAGACCTCCCCCACTGGGCCGCAAAACAAGTCTTCCTGAACATGGGTACACTGCTGCTCGGAGCCAGTGCGCTCGGCGTTGATGCCGTTCCCATGGAGGGAATGGACTTCAACACTCTGGATACGGAGTTTGGTTTGTCGGCAAAGGGACTCGCCCCAGTAGCGGTCGTAAGCCTGGGTTATCGGAAAGAGGACGATTTCAATGCCGCGTTGCCCAAGAGTAGGCTCCCCGAGGAACAGATTATGACTGTTATATAAGCCCGACTCTAGATCATAATCAGCCATACCACATTTGTAGAGGTTTTACTTGCTTTAAAAAACCTCCCACACCTCCCCCTGAACCTGAAACATAAAATGAATGCAATTGTTGTTGTTAACTTGTTTATTGCAGCTTATAATGGTTACAAATAAAGCAATAGCATCACAAATTTCACAAATAAAGCATTTTTTTCACTGCATTCTAGTTGTGGTTTGCCCAAACTCATCAATGTATCTTAGCGGCTCGAGGGTACCTCTAGAGATCCACTAGTGTCGACGATGTAGGTCACGGTCTCGAAGCCGCGGTGCGGGTGCCAGGGCGTGCCCTTGGGCTCCCCGGGCGCGTACTCCACCTCACCCATCTGGTCCATCATGATGAACGGGTCGAGGTGGCGGTAGTTGATCCCGGCGAACGCGCGGCGCACCGGGAAGCCCTCGCCCTCGAAACCGCTGGGCGCGGTGGTCACGGTGAGCACGGGACGTGCGACGGCGTCGGCGGGTGCGGATACGCGGGGCAGCGTCAGCGGGTTCTCGACGGTCACGGCGGGCATGTCGACACTAGTTCTAGCCAGCTTTTGTTCCCTTTAGTGAGGGTTAATTTCGAGCTTGGCGTAATCATGGTCATAGCTGTTTCCTGTGTGAAATTGTTATCCGCTCACAATTCCACACAACATACGAGCCGGAAGCATAAAGTGTAAAGCCTGGGGTGCCTAATGAGTGAGCTAACTCACATTAATTGCGTTGCGCTCACTGCCCGCTTTCCAGTCGGGAAACCTGTCGTGCCAGCTGCATTAATGAATCGGCCAACGCGCGGGGAGAGGCGGTTTGCGTATTGGGCGCTCTTCCGCTTCCTCGCTCACTGACTCGCTGCGCTCGGTCGTTCGGCTGCGGCGAGCGGTATCAGCTCACTCAAAGGCGGTAATACGGTTATCCACAGAATCAGGGGATAACGCAGGAAAGAACATGTGAGCAAAAGGCCAGCAAAAGGCCAGGAACCGTAAAAAGGCCGCGTTGCTGGCGTTTTTCCATAGGCTCCGCCCCCCTGACGAGCATCACAAAAATCGACGCTCAAGTCAGAGGTGGCGAAACCCGACAGGACTATAAAGATACCAGGCGTTTCCCCCTGGAAGCTCCCTCGTGCGCTCTCCTGTTCCGACCCTGCCGCTTACCGGATACCTGTCCGCCTTTCTCCCTTCGGGAAGCGTGGCGCTTTCTCATAGCTCACGCTGTAGGTATCTCAGTTCGGTGTAGGTCGTTCGCTCCAAGCTGGGCTGTGTGCACGAACCCCCCGTTCAGCCCGACCGCTGCGCCTTATCCGGTAACTATCGTCTTGAGTCCAACCCGGTAAGACACGACTTATCGCCACTGGCAGCAGCCACTGGTAACAGGATTAGCAGAGCGAGGTATGTAGGCGGTGCTACAGAGTTCTTGAAGTGGTGGCCTAACTACGGCTACACTAGAAGAACAGTATTTGGTATCTGCGCTCTGCTGAAGCCAGTTACCTTCGGAAAAAGAGTTGGTAGCTCTTGATCCGGCAAACAAACCACCGCTGGTAGCGGTGGTTTTTTTGTTTGCAAGCAGCAGATTACGCGCAGAAAAAAAGGATCTCAAGAAGATCCTTTGATCTTTTCTACGGGGTCTGACGCTCAGTGGAACGAAAACTCACGTTAAGGGATTTTGGTCATGAGATTATCAAAAAGGATCTTCACCTAGATCCTTTTAAATTAAAAATGAAGTTTTAAATCAATCTAAAGTATATATGAGTAAACTTGGTCTGACAGTTACCAATGCTTAATCAGTGAGGCACCTATCTCAGCGATCTGTCTATTTCGTTCATCCATAGTTGCCTGACTCCCCGTCGTGTAGATAACTACGATACGGGAGGGCTTACCATCTGGCCCCAGTGCTGCAATGATACCGCGAGACCCACGCTCACCGGCTCCAGATTTATCAGCAATAAACCAGCCAGCCGGAAGGGCCGAGCGCAGAAGTGGTCCTGCAACTTTATCCGCCTCCATCCAGTCTATTAATTGTTGCCGGGAAGCTAGAGTAAGTAGTTCGCCAGTTAATAGTTTGCGCAACGTTGTTGCCATTGCTACAGGCATCGTGGTGTCACGCTCGTCGTTTGGTATGGCTTCATTCAGCTCCGGTTCCCAACGATCAAGGCGAGTTACATGATCCCCCATGTTGTGCAAAAAAGCGGTTAGCTCCTTCGGTCCTCCGATCGTTGTCAGAAGTAAGTTGGCCGCAGTGTTATCACTCATGGTTATGGCAGCACTGCATAATTCTCTTACTGTCATGCCATCCGTAAGATGCTTTTCTGTGACTGGTGAGTACTCAACCAAGTCATTCTGAGAATAGTGTATGCGGCGACCGAGTTGCTCTTGCCCGGCGTCAATACGGGATAATACCGCGCCACATAGCAGAACTTTAAAAGTGCTCATCATTGGAAAACGTTCTTCGGGGCGAAAACTCTCAAGGATCTTACCGCTGTTGAGATCCAGTTCGATGTAACCCACTCGTGCACCCAACTGATCTTCAGCATCTTTTACTTTCACCAGCGTTTCTGGGTGAGCAAAAACAGGAAGGCAAAATGCCGCAAAAAAGGGAATAAGGGCGACACGGAAATGTTGAATACTCATACTCTTCCTTTTTCAATATTATTGAAGCATTTATCAGGGTTATTGTCTCATGAGCGGATACATATTTGAATGTATTTAGAAAAATAAACAAATAGGGGTTCCGCGCACATTTCCTCCGAAAAGTGCCACCTAAATTGTAAGCGTTAATATTTTTGTTAAAATTCGCGTTAAATTTTTTGTTAAATCAGCTCATTTTTTTAACCAATAGGCCGAAATCGGCAAAAATCCCCTTATAAATCAAAAAGAATAGACCGAGATAGGGTTGAGTGTTGTTCCCAGTTTTGGC |
| pKT111.XenA | AAAATAATCAATTAGGGGTTCCGCGCAACAATTTCCCCGAAAAAAGTGCCCACCTAAATTGTAAAGCGTTTAATATTTTTGTTTAAAATTCGCGTTAAATTTTTTGTTAAATCAGCTCATTTTTTTAACCAATAGGCCGAAAATCGGCAAAAATCCCTTATAAATCAAAAAGAATAGACCGAGATAGGGTTGAGTGTTGTTCCAGTTTGGAACAAGAGTCCACTATTAAAGAACGTGGACTCCAACGTCAAAGGGCGAAAAACCGTCTATCAGGGCGATGGCCCACTACGTGAACCATCACCCTAATCAAGTTTTTTGGGGTCGAGGTGCCGTAAAGCACTAAATCGGAACCCTAAAGGGAGCCCCCGATTTAGAGCTTGACGGGGAAAGCCGGCGAACGTGGCGAGAAAGGAAGGGAAGAAAGCGAAAGGAGCGGGCGCTAGGGCGCTGGCAAGTGTAGCGGTCACGCTGCGCGTAACCACCACACCCGCCGCGCTTAATGCGCCGCTACAGGGCGCGTCCCATTCGCCATTCAGGCTGCGCAACTGTTGGGAAGGGCGATCGGTGCGGGCCTCTTCGCTATTACGCCAGCTGGCGAAAGGGGGATGTGCTGCAAGGCGATTAAGTTGGGTAACGCCAGGGTTTTCCCAGTCACGACGTTGTAAAACGACGGCCAGTGAATTGTAATACGACTCACTATAGGGCGAATTGGGTACGTACCGGGCCCCTAGTATGTATGTAAGTTAATAAAACCCATTTTTGCGGAAAGTAGATAAAAAAAACATTTTTTTTTTTTACTGCACTGGATATCATTGAACTTATCTGATCAGTTTTAAATTTACTTCGATCCAAGGGTATTTGATGTACCAGGTTCTTTCGATTACCTCTCACTCAAAATGACATTCCACTCAAAGTCAGCGCTGTTTGCCTCCTTCTCTGTCCACAGAAATATCGCCGTCTCTTTCGCCGCTGCGTCCGCTATCTCTTTCGCCACCGTTTGTAGCGTTACGTAGCGTCAATGTCCGCCTTCAGTTGCATTTTGTCAGCGGTTTCGTGACGAAGCTCCAAGCGGTTTACGCCATCAATTAAACACAAAGTGCTGTGCCAAAACTCCTCTCGCTTCTTATTTTTGTTTGTTTTTTGAGTGATTGGGGTGGTGATTGGTTTTGGGTGGGTAAGCAGGGGAAAGTGTGAAAAATCCCGGCAATGGGCCAAGAGGATCAGGAGCTATTAATTCGCGGAGGCAGCAAACACCCATCTGCCGAGCATCTGAACAATGTGAGTAGTACATGTGCATACATCTTAAGTTCACTTGATCTATAGGAACTGCGATTGCAACATCAAATTGTCTGCGGCGTGAGAACTGCGACCCACAAAAATCCCAAACCGCAATTGCACAAACAAATAGTGACACGAAACAGATTATTCTGGTAGCTGTTCTCGCTATATAAGACAATTTTTGAGATCATATCATGATCAAGACATCTAAAGGCATTCATTTTCGACTATATTCTTTTTTACAAAAAATATAACAACCAGATATTTTAAGCTGATCCTAGATGCACAAAAAATAAATAAAAGTATAAACCTACTTCGTAGGATACTTCGGGGTACTTTTTGTTCGGGGTTAGATGAGCATAACGCTTGTAGTTGATATTTGAGATCCCCTATCATTGCAGGGTGACAGCGGAGCGGCTTCGCAGAGCTGCATTAACCAGGGCTTCGGGCAGGCCAAAAACTACGGCACGCTCCGGCCACCCAGTCCGCCGGAGGACTCCGGTTCAGGGAGCGGCCAACTAGCCGAGAACCTCACCTATGCCTGGCACAATATGGACATCTTTGGGGCGGTCAATCAGCCGGGCTCCGGATGGCGGCAGCTGGTCAACCGGACACGCGGACTATTCTGCAACGAGCGACACATACCGGCGCCCAGGAAACATTTGCTCAAGAACGGTGAGTTTCTATTCGCAGTCGGCTGATCTGTGTGAAATCTTAATAAAGGGTCCAATTACCAATTTGAAACTCAGTTTGCGGCGTGGCCTATCCGGGCGAACTTTTGGCCGTGATGGGCAGTTCCGGTGCCGGAAAGACGACCCTGCTGAATGCCCTTGCCTTTCGATCGCCGCAGGGCATCCAAGTATCGCCATCCGGGATGCGACTGCTCAATGGCCAACCTGTGGACGCCAAGGAGATGCAGGCCAGGTGCGCCTATGTCCAGCAGGATGACCTCTTTATCGGCTCCCTAACGGCCAGGGAACACCTGATTTTCCAAGCCATGGTGCGGATGCCACGACATCTGACCTATCGGCAGCGAGTGGCCCGCGTGGATCAGGTGATCCAGGAGCTTTCGCTCAGCAAATGTCAGCACACGATCATCGGTGTGCCCGGCAGGGTGAAAGGTCTGTCCGGCGGAGAAAGGAAGCGTCTGGCATTCGCCTCCGAGGCTCTAACCGATCCGCCGCTTCTGATCTGCGATGAGCCCACCTCCGGACTGGACTCCTTTACCGCCCACAGCGTCGTCCAGGTGCTGAAGAAGCTGTCGCAGAAGGGCAAGACCGTCATCCTGACCATTCATCAGCCGTCTTCCGAGCTGTTTGAGCTCTTTGACAAGATCCTTCTGATGGCCGAGGGCAGGGTAGCTTTCTTGGGCACTCCCAGCGAAGCCGTCGACTTCTTTTCCTAGTGAGTTCGATGTGTTTATTAAGGGTATCTAGTATTACATAACATCTCAACTCCTATCCAGCGTGGGTGCCCAGTGTCCTACCAACTACAATCCGGCGGACTTTTACGTACAGGTGTTGGCCGTTGTGCCCGGACGGGAGATCGAGTCCCGTGATCGGATCGCCAAGATATGCGACAATTTTGCCATTAGCAAAGTAGCCCGGGATATGGAGCAGTTGTTGGCCACCAAAAATCTGGAGAAGCCACTGGAGCAGCCGGAGAATGGGTACACCTACAAGGCCACCTGGTTCATGCAGTTCCGGGCGGTCCTGTGGCGATCCTGGCTGTCGGTGCTCAAGGAACCACTCCTCGTAAAAGTGCGACTTATTCAGACAACGGTGAGTGGTTCCAGTGGAAACAAATGATATAACGCTTACAATTCTTGGAAACAAATTCGCTAGATTTTAGATAGAATTGCCTGATTCCACACCCTTCTTAGTTTTTTTCAATGAGATGTATAGTTTATAGTTTTGCAGAAGATAAATAAATTTCATTTAACTCGCGAATATTAATGAGATGCGAGTAACATTTTAATTTGCAGATGGTTGCCATCTTGATTGGCCTCATCTTTTTGGGCCAACAACTCACGCAAGTGGGTGTGATGAATATCAACGGAGCCATCTTCCTCTTCCTGACCAACATGACCTTTCAAAACGTCTTTGCCACGATAAATGTAAGTCATGTTTAGAATACATTTGCATTTCAATAATTTACTAACTTTCTAATGAATCGATTCGATTTAGGTGTTCACCTCAGAGCTGCCAGTTTTTATGAGGGAGGCCCGAAGTCGACTTTATCGCTGTGACACATACTTTCTGGGCAAAACGATTGCCGAATTGCCGCTTTTTCTCACAGTGCCACTGGTCTTCACGGCGATTGCCTATCCGATGATCGGACTGCGGGCCGGAGTGCTGCACTTCTTCAACTGCCTGGCGCTGGTCACTCTGGTGGCCAATGTGTCAACGTCCTTCGGATATCTAATATCCTGCGCCAGCTCCTCGACCTCGATGGCGCTGTCTGTGGGTCCGCCGGTTATCATACCATTCCTGCTCTTTGGCGGCTTCTTCTTGAACTCGGGCTCGGTGCCAGTATACCTCAAATGGTTGTCGTACCTCTCATGGTTCCGTTACGCCAACGAGGGTCTGCTGATTAACCAATGGGCGGACGTGGAGCCGGGCGAAATTAGCTGCACATCGTCGAACACCACGTGCCCCAGTTCGGGCAAGGTCATCCTGGAGACGCTTAACTTCTCCGCCGCCGATCTGCCGCTGGACTACGTGGGTCTGGCCATTCTCATCGTGAGCTTCCGGGTGCTCGCATATCTGGCTCTAAGACTTCGGGCCCGACGCAAGGAGTAGCCGACATATATCCGAAATAACTGCTTGTTTTTTTTTTTTACCATTATTACCATCGTGTTTACTGTTTATTGCCCCCTCAAAAAGCTAATGTAATTATATTTGTGCCAATAAAAACAAGATATGACCTATAGAATACAAGTATTTCCCCTTCGAACATCCCCACAAGTAGACTTTGGATTTGTCTTCTAACCAAAAGACTTACACACCTGCATACCTTACATCAAAAACTCGTTTATCGCTACATAAAACACCGGGATATATTTTTTATATACATACTTTTCAAATCGCGCGCCCTCTTCATAATTCACCTCCACCACACCACGTTTCGTAGTTGCTCTTTCGCTGTCTCCCACCCGCTCTCCGCAACACATTCACCTTTTGTTCGACGACCTTGGAGCGACTGTCGTTAGTTCCGCGCGATTCGGTTCGCTCAAATGGTTCCGAGTGGTTCATTTCGTCTCAATAGAAATTAGTAATAAATATTTGTATGTACAATTTATTTGCTCCAATATATTTGTATATATTTCCCTCACAGCTATATTTATTCTAATTTAATATTATGACTTTTTAAGGTAATTTTTTGTGACCTGTTCGGAGTGATTAGCGTTACAATTTGAACTGAAAGTGACATCCAGTGTTTGTTCCTTGTGTAGATGCATCTCAAAAAAATGGTGGGCATAATAGTGTTGTTTATATATATCAAAAATAACAACTATAATAATAAGAATACATTTAATTTAGAAAATGCTTGGATTTCACTGGAACTAGGCTAGCATAACTTCGTATAATGTATGCTATACGAAGTTATGCTAGCGGATCCGGGAATTGGGAATTCACGTAAGTACTGTCTGCAGCGTAAGCTTCGTACGTAGCGACCGTCTCAAAGTACTGCCTTTCTGCGTTGGAAAACATCGCCTTTTTCGTCCAAAAGGAGTCCCCAGGTTCGATCCGCATGGCGTTGTGCGTGCGTGCCTTTCTTTTCAAATGATTACGGCTATTAACTTGGGGGCGTTAAGTTGGAAACACGTAAATTGCAGACTGCGATTAGAGTGACCATGAGTAGGAGTTCAAAATCTCCTGACATCATTTTCTTAAAACCTGCTTTGTTTTTTACATTTCTATTTAATATAACTCCTATTTGAATAAAAAAACAAAACAAGTTTAGATGTTAAGATATTAACTACATCCTTTGCTCCAAAGGGAGAGGGGAAGTTATGGAGTTAATTAATTTGCTGTTGGAAATCAATATGGAGTCAGAAATATAATGATTTACTAAACCTTATTGAATCGGTAACGATGCGAATTTATATTAAAATAGCTTTTATGAAACATTCAACAAAAATATTATTAATGTTGGCCCACTTTAGCAACCGGTTAGGTCTACCGGTTGGGCAAGCAAAGATTCACGCCCTGGTTCGAGTCCCAACTAGTCCTGCAAAATACCGCAGCAAGTTTTAGAGAGACCAAGTGCCATTACCTCTCCCACTTCAGTTATCGGTTATGCGGCGTTTAAGTCGACAGCTTGCCGTCTCTAGCTCCGGTGCCTATATAAAGCAGCCCGCTTTCCACATTTCATATTCGTTTTACGTTTGTCAAGCCTCATAGCCGGCAGTTCGAACGTATACGCTCTCTGAGTCAGACCTCGAAATCGTAGCTCTACACAATTCTGTGAATTTTCCTTGTCGCGTGTGAAACACTTCCAAGGCAATCTTACAAAATGAGTGCTTTGTTCGAGCCTTACACGTTGAAGGATGTTACACTGCGGAACCGAATTGCTATTCCACCCATGTGCCAGTACATGGCCGAAGATGGTTTGATAAATGACTGGCACCAAGTACATTACGCTTCCATGGCTCGGGGAGGTGCGGGTCTGCTCGTAGTAGAAGCAACGGCAGTAGCGCCCGAGGGAAGGATCACACCTGGCTGTGCTGGAATATGGAGTGATGCTCATGCTCAGGCGTTTGTACCAGTGGTGCAAGCGATAAAAGCAGCCGGCAGTGTGCCAGGCATACAGATTGCGCATGCGGGACGTAAGGCTAGCGCCAATCGACCCTGGGAAGGCGACGATCACATTGGTGCTGACGACGCTCGGGGCTGGGAAACAATCGCCCCATCCGCAATTGCATTCGGAGCTCATCTCCCAAATGTCCCCCGGGCCATGACACTGGATGACATCGCTCGCGTTAAACAGGACTTCGTGGACGCGGCACGTAGGGCGAGGGACGCTGGATTTGAGTGGATCGAATTGCATTTCGCACACGGCTACCTGGGTCAATCGTTTTTTTCGGAACACTCGAACAAACGCACCGACGCCTACGGCGGCTCGTTCGACAACCGAAGCCGTTTTCTGCTGGAAACACTCGCAGCCGTCAGGGAAGTGTGGCCCGAAAACTTGCCGCTGACGGCACGATTTGGAGTGCTCGAATATGATGGACGGGATGAGCAAACGTTGGAAGAGAGCATCGAACTGGCAAGGCGCTTCAAAGCCGGTGGTCTGGATTTGCTCTCCGTGTCGGTAGGTTTTACCATTCCTGAAACCAATATACCTTGGGGACCTGCTTTCATGGGTCCTATAGCAGAGAGGGTCAGGAGGGAAGCGAAACTCCCTGTGACCAGTGCATGGGGCTTCGGCACCCCCCAATTGGCTGAAGCCGCCCTCCAAGCAAACCAATTGGATTTGGTCAGTGTTGGCCGAGCCCACTTGGCCGACCCTCATTGGGCATACTTCGCAGCGAAAGAACTGGGTGTTGAAAAAGCGAGTTGGACTCTGCCTGCGCCATACGCTCACTGGCTGGAACGCTACCGCTAAGCCCGACTCTAGATCATAATCAGCCATACCACATTTGTAGAGGTTTTACTTGCTTTAAAAAACCTCCCACACCTCCCCCTGAACCTGAAACATAAAATGAATGCAATTGTTGTTGTTAACTTGTTTATTGCAGCTTATAATGGTTACAAATAAAGCAATAGCATCACAAATTTCACAAATAAAGCATTTTTTTCACTGCATTCTAGTTGTGGTTTGCCCAAACTCATCAATGTATCTTAGCGGCTCGAGGGTACCTCTAGAGATCCACTAGTGTCGACGATGTAGGTCACGGTCTCGAAGCCGCGGTGCGGGTGCCAGGGCGTGCCCTTGGGCTCCCCGGGCGCGTACTCCACCTCACCCATCTGGTCCATCATGATGAACGGGTCGAGGTGGCGGTAGTTGATCCCGGCGAACGCGCGGCGCACCGGGAAGCCCTCGCCCTCGAAACCGCTGGGCGCGGTGGTCACGGTGAGCACGGGACGTGCGACGGCGTCGGCGGGTGCGGATACGCGGGGCAGCGTCAGCGGGTTCTCGACGGTCACGGCGGGCATGTCGACACTAGTTCTAGCCAGCTTTTGTTCCCTTTAGTGAGGGTTAATTTCGAGCTTGGCGTAATCATGGTCATAGCTGTTTCCTGTGTGAAATTGTTATCCGCTCACAATTCCACACAACATACGAGCCGGAAGCATAAAGTGTAAAGCCTGGGGTGCCTAATGAGTGAGCTAACTCACATTAATTGCGTTGCGCTCACTGCCCGCTTTCCAGTCGGGAAACCTGTCGTGCCAGCTGCATTAATGAATCGGCCAACGCGCGGGGAGAGGCGGTTTGCGTATTGGGCGCTCTTCCGCTTCCTCGCTCACTGACTCGCTGCGCTCGGTCGTTCGGCTGCGGCGAGCGGTATCAGCTCACTCAAAGGCGGTAATACGGTTATCCACAGAATCAGGGGATAACGCAGGAAAGAACATGTGAGCAAAAGGCCAGCAAAAGGCCAGGAACCGTAAAAAGGCCGCGTTGCTGGCGTTTTTCCATAGGCTCCGCCCCCCTGACGAGCATCACAAAAATCGACGCTCAAGTCAGAGGTGGCGAAACCCGACAGGACTATAAAGATACCAGGCGTTTCCCCCTGGAAGCTCCCTCGTGCGCTCTCCTGTTCCGACCCTGCCGCTTACCGGATACCTGTCCGCCTTTCTCCCTTCGGGAAGCGTGGCGCTTTCTCATAGCTCACGCTGTAGGTATCTCAGTTCGGTGTAGGTCGTTCGCTCCAAGCTGGGCTGTGTGCACGAACCCCCCGTTCAGCCCGACCGCTGCGCCTTATCCGGTAACTATCGTCTTGAGTCCAACCCGGTAAGACACGACTTATCGCCACTGGCAGCAGCCACTGGTAACAGGATTAGCAGAGCGAGGTATGTAGGCGGTGCTACAGAGTTCTTGAAGTGGTGGCCTAACTACGGCTACACTAGAAGAACAGTATTTGGTATCTGCGCTCTGCTGAAGCCAGTTACCTTCGGAAAAAGAGTTGGTAGCTCTTGATCCGGCAAACAAACCACCGCTGGTAGCGGTGGTTTTTTTGTTTGCAAGCAGCAGATTACGCGCAGAAAAAAAGGATCTCAAGAAGATCCTTTGATCTTTTCTACGGGGTCTGACGCTCAGTGGAACGAAAACTCACGTTAAGGGATTTTGGTCATGAGATTATCAAAAAGGATCTTCACCTAGATCCTTTTAAATTAAAAATGAAGTTTTAAATCAATCTAAAGTATATATGAGTAAACTTGGTCTGACAGTTACCAATGCTTAATCAGTGAGGCACCTATCTCAGCGATCTGTCTATTTCGTTCATCCATAGTTGCCTGACTCCCCGTCGTGTAGATAACTACGATACGGGAGGGCTTACCATCTGGCCCCAGTGCTGCAATGATACCGCGAGACCCACGCTCACCGGCTCCAGATTTATCAGCAATAAACCAGCCAGCCGGAAGGGCCGAGCGCAGAAGTGGTCCTGCAACTTTATCCGCCTCCATCCAGTCTATTAATTGTTGCCGGGAAGCTAGAGTAAGTAGTTCGCCAGTTAATAGTTTGCGCAACGTTGTTGCCATTGCTACAGGCATCGTGGTGTCACGCTCGTCGTTTGGTATGGCTTCATTCAGCTCCGGTTCCCAACGATCAAGGCGAGTTACATGATCCCCCATGTTGTGCAAAAAAGCGGTTAGCTCCTTCGGTCCTCCGATCGTTGTCAGAAGTAAGTTGGCCGCAGTGTTATCACTCATGGTTATGGCAGCACTGCATAATTCTCTTACTGTCATGCCATCCGTAAGATGCTTTTCTGTGACTGGTGAGTACTCAACCAAGTCATTCTGAGAATAGTGTATGCGGCGACCGAGTTGCTCTTGCCCGGCGTCAATACGGGATAATACCGCGCCACATAGCAGAACTTTAAAAGTGCTCATCATTGGAAAACGTTCTTCGGGGCGAAAACTCTCAAGGATCTTACCGCTGTTGAGATCCAGTTCGATGTAACCCACTCGTGCACCCAACTGATCTTCAGCATCTTTTACTTTCACCAGCGTTTCTGGGTGAGCAAAAACAGGAAGGCAAAATGCCGCAAAAAAGGGAATAAGGGCGACACGGAAATGTTGAATACTCATACTCTTCCTTTTTCAATATTATTGAAGCATTTATCAGGGTTATTGTCTCATGAGCGGATACATATTTGAATGTATTTAGAAAAATAAACAAATAGGGGTTCCGCGCACATTTCCTCCGAAAAGTGCCACCTAAATTGTAAGCGTTAATATTTTTGTTAAAATTCGCGTTAAATTTTTTGTTAAATCAGCTCATTTTTTTAACCAATAGGCCGAAATCGGCAAAAATCCCCTTATAAATCAAAAAGAATAGACCGAGATAGGGTTGAGTGTTGTTCCCAGTTTTGGC |

**Supplementary references**

1.	Venken, K. J. T., He, Y., Hoskins, R. A. & Bellen, H. J. P[acman]: A BAC transgenic platform for targeted insertion of large DNA fragments in *D. melanogaster*. *Science (1979)* **314**, 1747–1751 (2006).

2.	Hazelrigg, T., Levis, R. & Rubin, G. M. Transformation of *white* locus DNA in Drosophila: Dosage compensation, *zeste* interaction, and position effects. *Cell* **36**, 469–481 (1984).

3.	Clark, M. *et al.* Bioremediation of Industrial Pollutants by Insects Expressing a Fungal Laccase. *ACS Synth Biol* **11**, 308–316 (2022).
